# Tonically active GABAergic neurons in the dorsal periaqueductal gray control the initiation and execution of instinctive escape

**DOI:** 10.1101/2023.11.03.565561

**Authors:** A. Vanessa Stempel, Dominic A. Evans, Oriol Pavón Arocas, Federico Claudi, Stephen C. Lenzi, Elena Kutsarova, Troy W. Margrie, Tiago Branco

## Abstract

To avoid predation, animals perform defensive actions that are both instinctive and adaptable to the environment. In mice, the decision to escape from imminent threats is implemented by a feed-forward circuit in the midbrain, where excitatory VGluT2^+^ neurons in the dorsal periaqueductal gray (dPAG) compute escape initiation and escape vigour from threat evidence. Here we show that GABAergic VGAT^+^ neurons in the dPAG dynamically control this process by modulating the excitability of excitatory escape neurons. Using *in vitro* patchclamp and *in vivo* neural activity recordings in freely behaving mice we found that VGAT^+^ dPAG neurons fire action potentials tonically in the absence of synaptic inputs and are a major source of synaptic inhibition to VGluT2^+^ dPAG neurons. Activity in these spontaneously firing VGAT^+^ cells transiently decreases at escape onset and increases during escape, peaking at escape termination. Optogenetically increasing or decreasing VGAT^+^ dPAG activity bidirectionally changes the probability of escape when the stimulation is delivered at the time of threat onset, and the duration of escape when delivered after escape initiation. We conclude that the activity of tonically firing VGAT^+^ dPAG neurons sets a threshold for escape initiation and controls the execution of the flight locomotor action.

## Introduction

Escape behaviour is a set of locomotor actions that move an animal away from threat. While these actions can be reflexive and stereotyped to ensure short reaction times and fast movements, it is equally important for survival that they are flexible (1–3). For example, the probability of initiating escape depends on predation risk and threat history (4–11), as well as on the internal state (12–16). In environments where predators abound, it is more likely that ambiguous sensory stimuli might signal the presence of predators and thus the probability of initiating escape should increase (17). Likewise, when faced with a competing motivation, such as hunger, the probability of defensive behaviour decreases (12, 18).

Beyond flexibility in escape initiation, it is equally important that the execution of escape is flexible. The flight to safety should not be ballistic (i.e., along a pre-determined trajectory with fixed kinetics) but instead should allow for continuous adjustments of vigour and trajectory to deal with variability in the terrain, adjust escape to unexpected events and terminate escape at the appropriate place and time (19– 22). Accordingly, recent work has shown that mice on a fast shelter-directed escape trajectory can slow down and navigate around an obstacle that suddenly appears, while maintaining the goal of reaching safety (20). This degree of flexibility in escape initiation and execution suggests the presence of modulatory components acting on the neural circuits controlling instinctive escape. In particular, the ability of decreasing the sensitivity to threat and slowing down locomotion during escape suggests the presence of a regulatory inhibitory network.

In mammals, the key circuit for commanding escape behaviour is the periaqueductal gray (PAG), located in the midbrain (23–25). The PAG is an interface between the forebrain and central pattern generators in the hindbrain and spinal cord that acts as a critical neural circuit node for integrating sensory and state-dependent information, as well as for executing instinctive behaviours (25–39). In the dorsal PAG (dPAG), excitatory VGluT2^+^ neurons receive information about imminent visual and auditory threats from the superior colliculus through a feed-forward excitatory connection, and their activation determines the onset of escape and controls escape vigour (31). While modulation of threat processing and escape behaviour can be accomplished through a variety of different circuits (40–46), the network of excitatory neurons in the dPAG stands out as a key control node (24, 29, 31, 47–50). Previous work has demonstrated the presence of an inhibitory tone in the PAG (51–53) that could in principle control the excitability of the excitatory network to modulate escape. For example, local application of a GABA_A_ receptor antagonist in the PAG elicits responses to a previously sub-threshold stimulus (23, 24). Disinhibition of the PAG has been suggested as a mechanism for initiating other instinctive behaviours, such as vocalisations and freezing (51, 54), with inhibitory long-range projections from forebrain regions inhibiting local GABAergic circuits within the PAG. Also, both *in vivo* and *in vitro* single-unit recordings from GABAergic neurons in the lateral and ventral PAG have shown that their baseline firing rates are higher than excitatory neurons (35, 51–54). Here we have tested the hypothesis that local GABAergic neurons within the dPAG control escape behaviour by setting the excitability of the dPAG escape network.

## Results

### GABAergic neurons in dPAG fire tonically in the absence of synaptic input

To investigate how inhibitory neurons in the dPAG might control the excitability of the local excitatory neurons, we performed patch-clamp recordings from identified VGAT^+^ and VGluT2^+^ neurons in acute midbrain slices *in vitro*. While whole-cell recordings of VGAT^+^ dPAG neurons showed biophysical and synaptic input properties similar to those of VGluT2^+^ dPAG cells (Figure S1, for comparison to VGluT2^+^ dPAG neurons, see (31)), recordings in loose-seal cell-attached configuration to monitor spiking non-invasively showed a marked difference between these two cell populations. VGAT^+^ neurons fired action potentials spontaneously at a mean firing rate of 4.7 ± 0.70 Hz, which was not significantly altered after pharmacological block of glutamatergic and inhibitory synaptic transmission (mean firing rate = 6.2 ± 0.84 Hz; P = 0.21, Mann-Whitney test between control and blockers; Figure 1A; see Figure S2 for anatomical locations). Using inter-spike interval (ISI) analysis and a measure of variability based on the coefficient of variation (CV_2_) (55), we found the firing of VGAT^+^ dPAG neurons to be highly regular, as expected from a dominant cell intrinsic mechanism (CV_2_ = 0.33 ± 0.03; Poisson firing statistics have a CV_2_ of 1 and a perfectly regular neuron has a CV_2_ of 0; Figure S3A, B). In contrast, the majority of VGluT2^+^ dPAG neurons were not spontaneously active both in control conditions (92.8 %, 39 out of 42 cells, firing rate < 0.04 Hz; mean firing rate across all neurons = 0.11 ± 0.068 Hz) or after block of inhibitory synaptic transmission (mean firing rate in picrotoxin = 0.039 ± 0.033 Hz; P = 0.85, Mann-Whitney tests between control and picrotoxin; Figure 1B). These data show that VGAT^+^ dPAG neurons are well positioned to provide sustained inhibition to the local network through synaptic input-independent action potential firing.

**Fig. 1.**
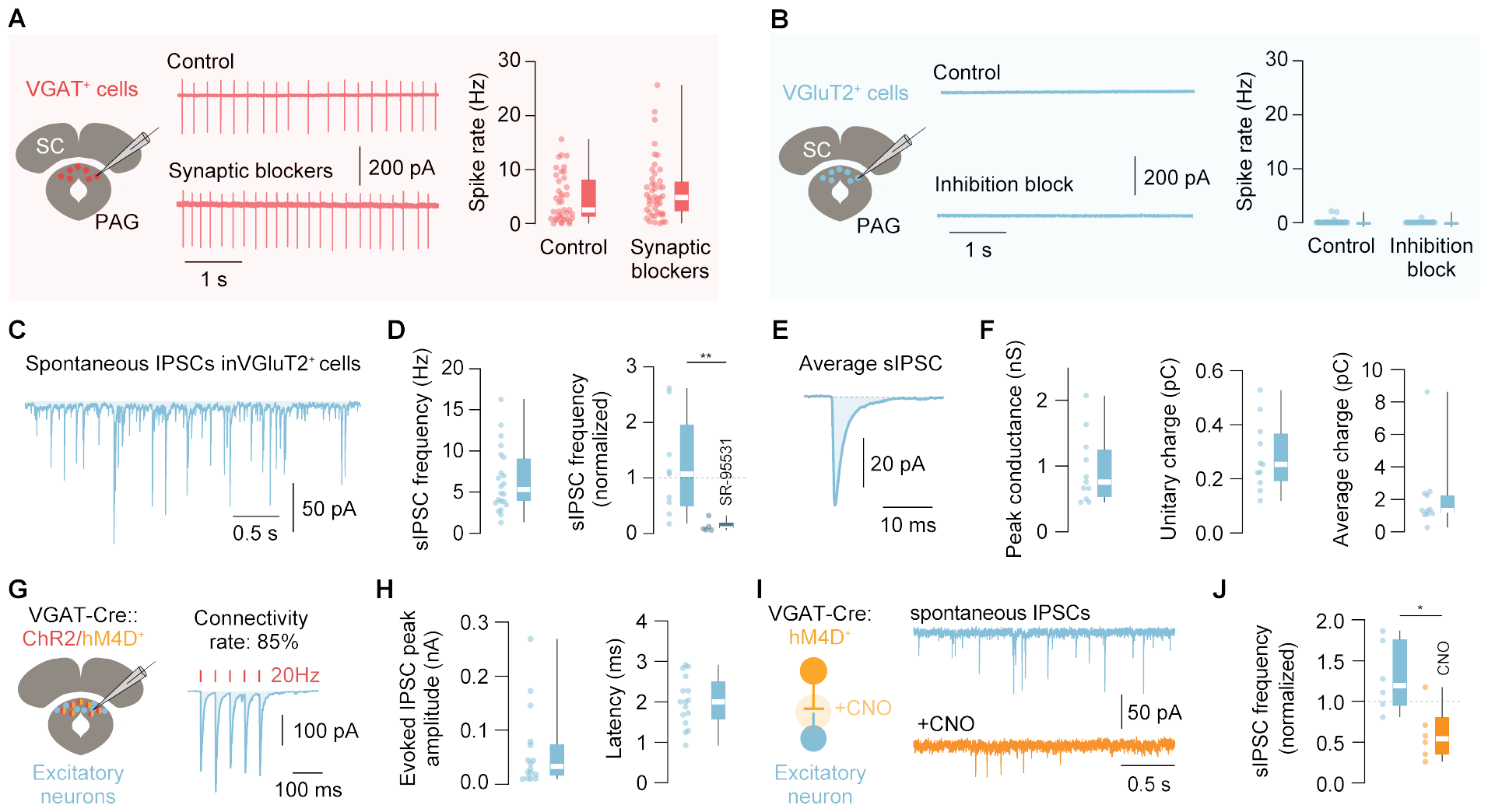
Properties of GABAergic inhibition in the dorsal PAG. **A**. Schematic of a patch-clamp recording from visually identified VGAT^+^ neurons in the dPAG (left). Loose-seal cell-attached recordings of VGAT^+^ dPAG neurons (middle): Example current traces of spontaneously spiking VGAT^+^ neurons without (control, top) and with synaptic blockers (2 mM kynurenic acid and 50 µM picrotoxin, bottom) added to the bath. Summary of VGAT^+^ dPAG neuron firing rate for control (n = 39 cells, N = 20 mice) and in synaptic blockers (n = 45 cells, N = 13 mice) (right panels). **B**. Same as A, for VGluT2^+^ dPAG neurons without blockers (control) and with a blocker of synaptic inhibition (50 µM picrotoxin). Summary of VGluT2^+^ dPAG neuron firing rate for control (n = 42 cells, N = 13 mice) and in 50 µM picrotoxin (n = 30 cells, N = 9). Comparison between VGAT^+^ and VGluT2^+^ controls: p<0.0001, Mann-Whitney test. **C**. Example whole-cell patch-clamp recording of spontaneous IPSCs onto a VGluT2^+^ dPAG neuron in the presence of 2 mM kynurenic acid at holding potential of -70mV. **D**. Summary graphs of the mean spontaneous IPSC frequency recorded in VGluT2^+^ dPAG neurons from n = 22, N = 8 mice (left panel) and of the normalized mean spontaneous IPSC frequency after bath application of SR-95331 (n = 5, N = 4 mice) and time-matched controls (n = 9, N = 7 mice) (right panel). **E**. Average spontaneous IPSC from the same neuron as in C. **F**. Summary graphs of the mean peak phasic conductance (left panel), unitary IPSC charge (middle panel) and average IPSC charge (right panel) from n = 11 cells, N = 2 mice. **G**. Schematic of the experiment for panels G–J (left). AAV-DIO-ChR2 and AAV-DIO-hM4D were co-expressed in the dPAG of VGAT-Cre animals and patch-clamp recordings were performed from presumptive, non-spiking excitatory neurons. Connectivity rate between ChR2-expressing VGAT^+^ neurons and presumptive excitatory dPAG neurons (n = 17 / 20 cells; N = 2 mice) with an example trace of inhibitory currents recorded in voltage-clamp configuration upon blue light stimulation (5 pulses, 1 ms duration, 20 Hz inter-stimulus interval) (right panel). **H**. Summary graphs of the evoked IPSC amplitude (left panel) and synaptic delay of this connection (right panel) (n = 16 cells, N = 2 mice). **I**. Same experiment as shown in G, with bath application of CNO (left panel). Example traces of spontaneous IPSCs recorded in 2 mM kynurenic acid before (blue trace) and after (orange trace) bath application of 10 µM CNO (right panel). **J**. Summary of the normalized change of spontaneous IPSC frequency in control conditions vs. CNO (n = 6 cells each, N = 2 mice). Box-and-whisker plots show median, IQR and range, and individual data points. See also Figures S1–S3.

### VGluT2+ neurons in dPAG are phasically inhibited by local GABAergic neurons

To assess the impact of local tonically active GABAergic neurons on dPAG excitability, we performed whole-cell recordings from VGluT2^+^ dPAG neurons *in vitro* and first isolated spontaneous inhibitory postsynaptic currents (IPSCs) by blocking glutamatergic synaptic transmission. We found frequent phasic inhibition (mean IPSC frequency = 6.5 ± 0.83 Hz) that was selectively blocked by a GABA_A_ receptor antagonist (SR-95531: mean fraction= 0.13 ± 0.05; control: mean fraction = 1.17 ± 0.3, P = 0.002, Mann-Whitney test, Figure 1C, D). Phasic inhibitory currents had a peak conductance of 0.94 ± 0.16 nS and a unitary IPSC charge of 0.28 ± 0.04 pC, generating a continuous average charge over time of 2.12 ± 0.68 pC (Figure 1E, F). These data show that VGluT2^+^ dPAG neurons receive sustained phasic GABA_A_ receptor-mediated inhibition with an inhibitory charge transfer that should in principle be large enough to regulate their input-output function *in vivo* (as e.g. shown for cerebellar granule cells that have a similarly high input resistance (56)).

Next, we tested the contribution of local VGAT^+^ dPAG neurons to inhibition of VGluT2^+^ dPAG cells. We used a combined optoand chemogenetic strategy to co-express an inhibitory DREADD (designer receptor exclusively activated by designer drugs, hM4Di-mCherry) and channelrhodopsin2 (ChR2-YFP) in the dPAG of VGAT-Cre mice to selectively and bidirectionally change the activity of VGAT^+^ dPAG neurons while recording inhibitory currents in excitatory dPAG cells. We first optogenetically activated VGAT^+^ dPAG neurons and found monosynaptic inhibitory inputs in 85% of glutamatergic dPAG neurons (mean latency = 2.0 ± 0.15 ms), with a mean evoked IPSC amplitude of -62.58 ± 18.31 pA and a facilitating paired pulse ratio (1.17 ± 0.16; Figure 1G, H). We then chemogenetically blocked synaptic transmission from VGAT^+^ dPAG neurons with Clozapine-N-Oxide (CNO, 10 µM), which reduced the frequency of spontaneous IPSCs onto glutamatergic neurons by 70 % in comparison to control recordings (normalized change in spontaneous IPSC frequency for CNO: mean = 0.59 ± 0.14; control: mean = 1.29 ± 0.17; P = 0.01, unpaired t-test with Welch’s correction; Figures 1I, J and S3A). Application of CNO also significantly reduced the amplitude of ChR2-evoked IPSCs (normalized peak amplitude CNO = 0.39 ± 0.16; control = 1.35 ± 0.33; CNO vs. control: P = 0.042, unpaired t-test; Figure S3C, D) further confirming the efficacy and specificity of our chemogenetic inhibition approach. It is important to note that since these experiments rely on small targeted injections and co-expression of two viral constructs within the same cells, our quantification of the contribution of local VGAT^+^ dPAG to spontaneous inhibition is likely to be an underestimate. Together these data show that VGAT^+^ neurons in the dPAG provide phasic and sustained monosynaptic GABA_A_ receptor-mediated inhibition onto VGluT2^+^ dPAG cells.

### VGAT+ dPAG neurons are spontaneously active in vivo and modulated during escape behaviour

We next measured the activity of VGAT^+^ dPAG neurons *in vivo* to determine whether they are a source of sustained inhibition as suggested by our *in vitro* experiments, and how their activity relates to escape behaviour. Given the characteristic firing profile of VGAT^+^ dPAG neurons *in vitro*, we first performed chronic silicon probe recordings from the dPAG of freely moving mice to determine whether there are single units with similar activity. Analysis of firing rates for periods when mice were spontaneously exploring the behavioural arena revealed the existence of neurons with high baseline firing rates across the recorded population (9/19 neurons with mean firing rates above 8 Hz; mean frequency = 32.2 ± 7.0 Hz; N = 3 mice; Figure S3E–G). This is consistent with previous reports of high baseline firing rates of VGAT^+^ neurons *in vivo* in other subdivisions of the PAG (35, 54), and with synaptic input driving neurons to higher firing rates than *in vitro*.

To directly record neural activity from molecularly-identified VGAT^+^ neurons, we performed *in vivo* calcium imaging of VGAT^+^ dPAG neurons expressing GCaMP6s in freely moving mice using a head-mounted miniature microscope (Figure S4). In line with the *in vitro* cell-attached and *in vivo* single-unit recordings during exploration, we found that VGAT^+^ dPAG neurons were active while naïve animals freely explored the arena (median calcium event rate = 0.1 Hz, IQR = [0.090, 0.11 Hz], 209 neurons, N = 8 mice; Figures 2A–C and S5A). To test whether this activity was correlated with the locomotor state of the animal, we calculated a locomotion modulation index for each neuron (57), and found that only a small fraction of cells was significantly modulated by movement (4.5 % of cells positively, 4.5 % of cells negatively modulated; Figure S5B). Together with the previous results, these data consolidate the view that VGAT^+^ dPAG neurons provide a sustained inhibitory tone on the dPAG network.

To understand how local VGAT^+^ dPAG activity might contribute to threat-evoked escape, we presented threatening overhead looming or ultrasound stimuli in the presence of a shelter (58–60). During stimulus-evoked escape to shelter, more than half of all active cells showed escape-related activity (median across mice = 54%, IQR = [46%, 76%], 123 neurons, N = 8 mice, Figures 2D, S5C–F). To compare the activity profiles of neurons across trials, where the escape sequence is stereotyped but has a variable duration, we linearly time warped each neuron’s activity to the median time-series of discrete events in the escape sequence to generate an average over trials (see Methods). Across the recorded population, the neuronal activity profiles were best described by two clusters identified by K-means clustering using scores from principal component analysis on the pooled mean time-warped responses (Figures 2E–H and S5D). Neurons belonging to both clusters were present in all animals (Figure S5E, F): the first cluster showed a gradual increase in activity from threat stimulus onset, peaking at escape termination (mean z-score between startle and escape stop = 0.69 ± 0.05 Z, n = 74 neurons, vs 0.06 ± 0.07 Z for same period in trials where stimulus failed to elicit escape, n = 41 neurons; two-tailed t-test, P = 1.1 x 10^-11^); Figure 2E–G); whereas the second cluster exhibited a transient decrease in activity with the trough coinciding with escape onset (mean z-score between startle and escape stop -0.43 ± 0.06 Z, n = 49 neurons, vs -0.13 ± 0.09 Z for same period in trials where animals failed to escape, n = 37 neurons; P = 0.006; 100 trials, N = 8 mice; 2E–H). To further test whether the activity of VGAT^+^ dPAG neurons was specifically modulated during escape, we compared activity profiles between fast spontaneous runs outside the threat stimulus context with slow escapes, where the average locomotor speed was lower than the spontaneous runs. We found that VGAT^+^ dPAG activity did not change during exploratory runs (*cluster 1, runs during exploration*, mean signal -0.12 ± 0.12 Z and -0.2 ± 0.13 Z in the 2 s before and after run onset respectively, two-tailed paired t-test, P = 0.66; n = 21 neurons, 66 trials, N = 4 mice) but changed significantly during slow escapes (*cluster 1, slow escapes*, mean signal -0.18 ± 0.20 Z before and +0.39 ± 0.17 Z after, two-tailed paired t-test, P = 0.047, n = 18 neurons, 15 trials, N = 4 mice; Figure S5G, H). In addition, cluster population activity was not correlated to the peak speed, acceleration or deceleration of the escape movement (Figure S6). Together this indicates that in the behavioural context tested, changes in the activity of VGAT^+^ dPAG neurons are highly specific for the escape action. These results suggest that VGAT^+^ dPAG neurons contribute to escape initiation by decreasing their activity in response to threat, thereby releasing the excitatory dPAG neurons from inhibition and facilitating their recruitment to initiate escape. In addition, the gradual increase in activity during escape indicates that VGAT^+^ dPAG neurons might also participate in shutting down the excitatory drive in the dPAG network to terminate the escape movement.

**Fig. 2.**
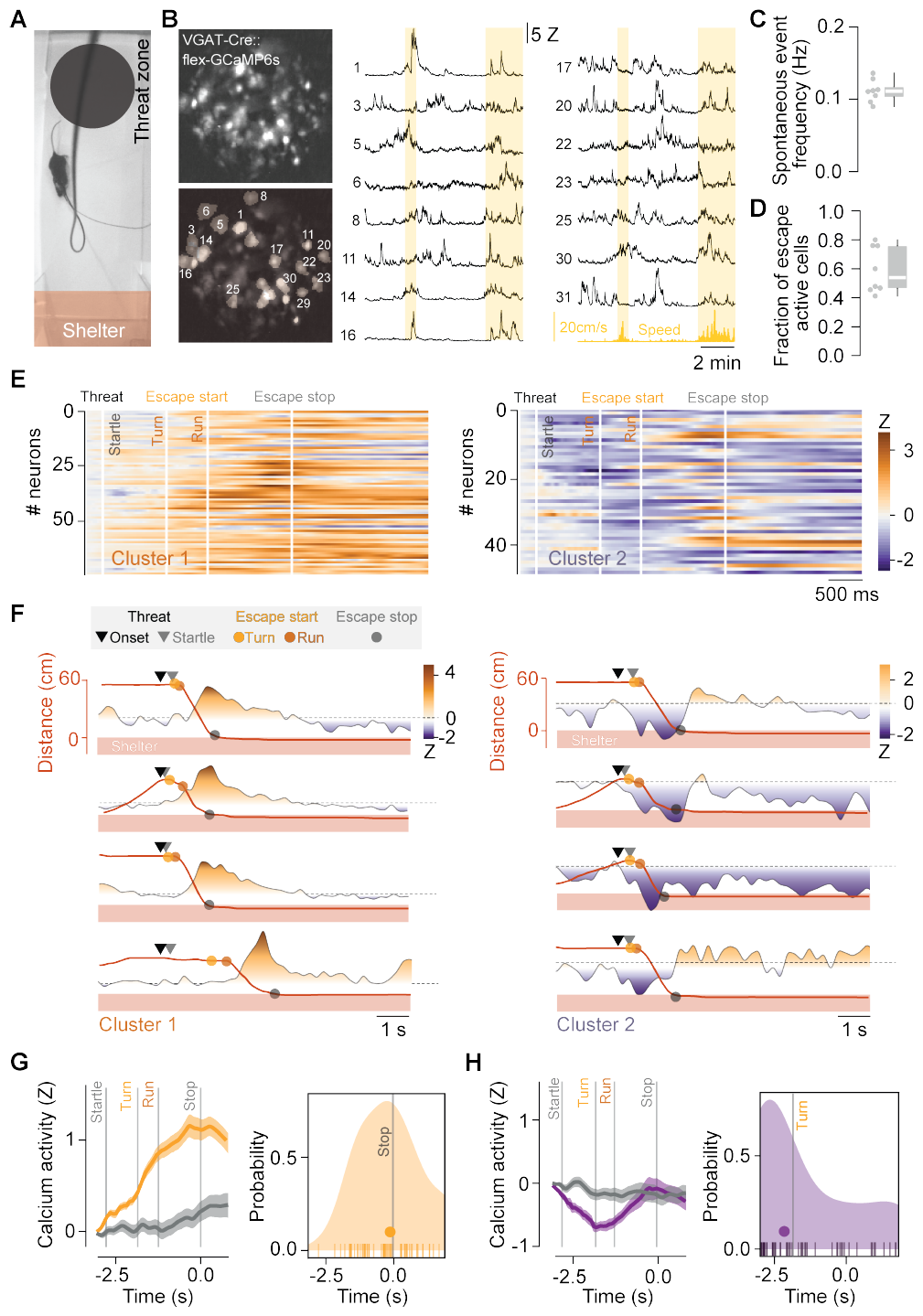
Calcium activity profiles of VGAT^+^ dPAG neurons during exploration and instinctive escape. **A**. Video frame during the escape assay showing the threat zone, threat (dark expanding disk) and shelter. **B**. Top left: example field-of-view (FOV) in the dPAG of a VGAT-Cre mouse expressing GCaMP6s. Bottom left: overlay of the same FOV with the extracted and numbered cell masks. Right: example traces of spontaneous calcium activity (Z-scored) of the indicated cell masks during baseline exploration. The speed of the animal is indicated in yellow and time windows of locomotor activity are shaded in light yellow. **C**. Summary of the mean spontaneous event frequency per animal. **D**. Fraction of cells active during escape. **E**. Heatmaps of the Z-scored calcium signal of all recorded neurons (mean calcium signal across trials) where each trial is time warped to the median temporal sequence of behavioural events of the cohort. Neurons are clustered on their activity from startle until escape stop (cluster 1, n = 74 neurons, left; cluster 2, n = 49 neurons, right; N = 8 mice). **F**. Activity of two example cell for cluster 1 (left) and cluster 2 (right) for 4 trials of the same recording session (rows). Calcium signal is coloured by Z-score and the animal’s distance from the shelter over time is shown in red. Markers show the onsets of the stimulus and behavioural events. All traces are aligned to stimulus onset without time warping. **G**. Left: average time warped signal of all cluster 1 neurons for trials where animals escaped (orange) and failed to escape (grey) to threat. Right: distribution of the time of peak mean calcium signal relative to escape termination (median time = -0.22 s, IQR = [-0.99, 0.56], filled circle; not different from escape stop = 0 s, *P* = 0.41, significantly different from startle, escape turn and run, *P* = 2.2x10^-31^, 1.1x10^-20^, 1.5x10^-12^ respectively, one-sample t-tests, Holm-Bonferroni corrected). **H**. Left: same as G for cluster 2. Right: distribution of the time of the minimum mean calcium signal relative to escape termination (median time of minimum signal = -2.18 s, IQR = [-2.93, -0.18], filled circle; not different from startle or escape turn, *P* = 0.19, 0.80 respectively, significantly different from escape run and stop, *P* = 0.01, 3.8 x10^-6^, sign tests, Holm-Bonferroni corrected). Box-and-whisker plots show median, IQR and range. See also Figures S4–S6.

### Activity manipulations of VGAT+ neurons bidirectionally modulate escape initiation and termination

The neural activity profiles of VGAT^+^ dPAG neurons recorded *in vivo* predict that: (1) manipulating their activity at the time of threat presentation should bidirectionally affect escape initiation, and (2) modulating activity during the escape action should perturb the duration of the escape action. We tested these predictions using bilateral, optogenetic manipulations of VGAT^+^ dPAG neurons during instinctive escape (activation with ChR2 and inactivation with iChloC, a chloride-conducting ChR; Figure S7A–C). First, we tested the effect of activity manipulations on escape initiation. Optogenetic activation of VGAT^+^ dPAG neurons at the time of threat presentation strongly reduced escape probability (mean escape probability = 0.88 ± 0.04 for threat only vs. 0.13 ± 0.05 for threat with laser on, P = 0.0005, paired two-tailed t-test; Figure 3A, C), but had no effect when it preceded the threat stimulus (end of the laser pulse train coincident with threat stimulus onset; mean escape probability = 0.92 ± 0.08 for threat with laser preceding stimulus onset vs. threat only, P = 0.44, Mann Whitney test; Figure S7F). Conversely, optogenetic inhibition of VGAT^+^ neurons during presentation of a weakly-threatening, low contrast stimulus, increased escape probability significantly (mean escape probability = 0.28 ± 0.06 for threat only, 0.78 ± 0.1 for threat with laser on; P = 0.0014, paired two-tailed t-test; Figure 3B, D). These results confirm that activity of VGAT^+^ dPAG neurons plays an important role in setting the escape threshold. We next tested the role of VGAT^+^ neurons in escape execution. Optogenetic stimulation of VGAT^+^ dPAG neurons with ChR2 after escape initiation often led to premature termination of escape, with the animal stopping before reaching the shelter (normalized mean escape termination distance from shelter, laser off = -0.10 ± 0.01, laser on = 0.09 ± 0.01; P < 0.001, paired two-tailed t-test; Figure 3E). Inactivation of VGAT^+^ neurons during escape had the opposite effect, with animals frequently overshooting their preferred target location of the arena shelter, resulting in a significant increase in total escape distance (laser off = 49.9 ± 4.6 cm, laser on = 69.4 ± 4.8 cm, P = 0.0097, unpaired t-test; Figure S8). Importantly, neither manipulation affected the mean speed during baseline exploratory locomotion (ChR2: P = 0.41, iChloc: P = 0.88; two-tailed paired t-tests; Figure S7D–H). These data show that the firing rate of VGAT^+^ dPAG neurons is a key variable that controls the execution of escape behaviour.

**Fig. 3.**
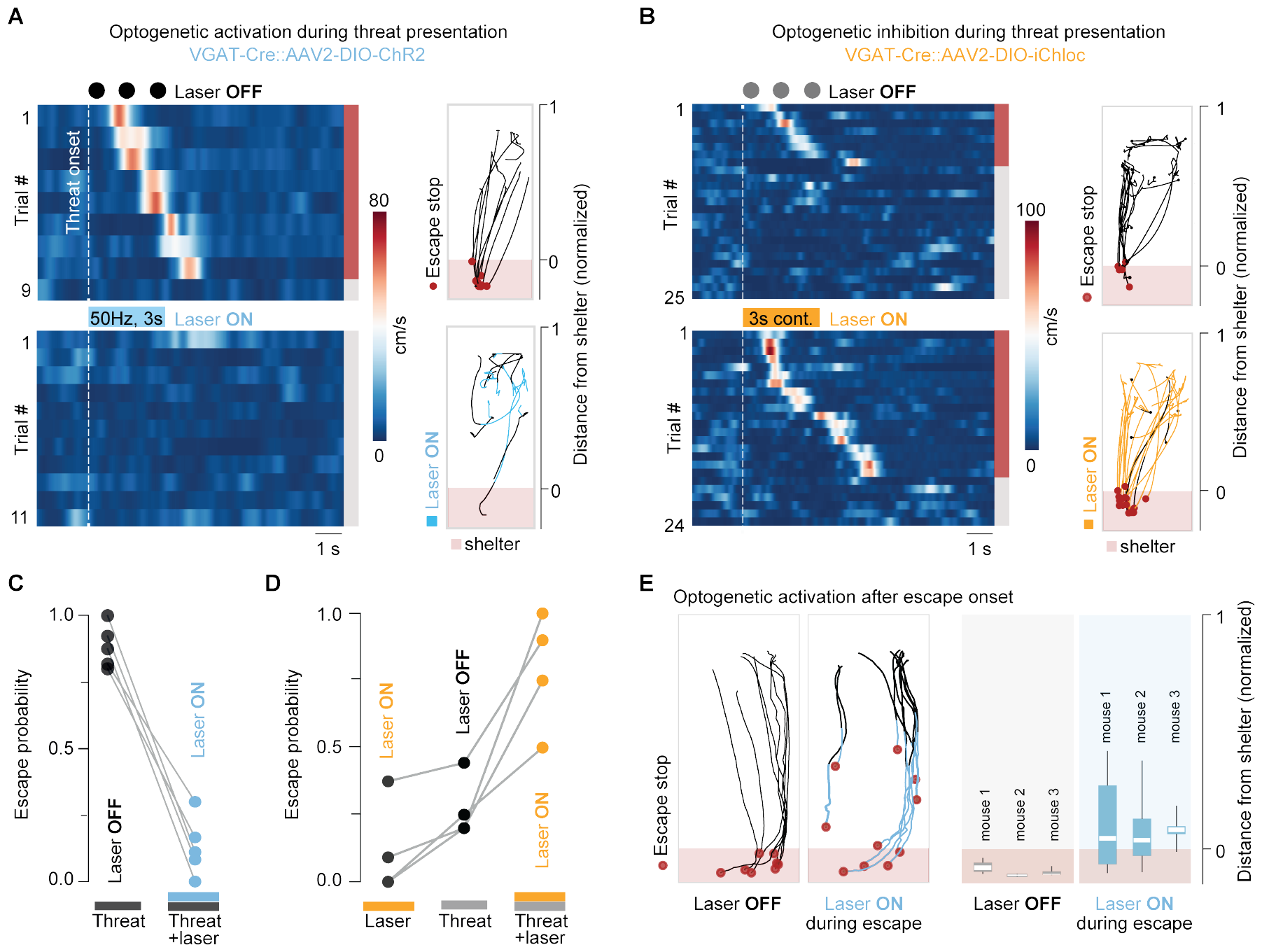
VGAT^+^ dPAG neurons bidirectionally modulate escape. **A**. Left: example raster plots of mouse speed for the same animal during a high-threat stimulus (a sequence of three high-contrast overhead looming spots) with 8 escapes in 9 trials (control, top) and 0 escapes in 11 trials during simultaneous optogenetic activation of the dPAG VGAT^+^ neurons (ChR2, 50 Hz, 5 s; bottom). Coloured bar indicates escape (red) and no escape (grey) trials. Laser OFF and Laser ON trials were interleaved during the experiment, and trials are sorted by reaction time. Right: Single trial mouse tracking of the same trials showing successful escapes to shelter (red dots = escape termination) for control (top) and during ChR2 stimulation (blue, bottom). **B**. Same as A, for optogenetic inactivation with iChloC (3–5 s, continuous). Left: example raster plots for the same animal during a low-threat stimulus (a sequence of three low-contrast looming spots) with 8 escapes in 25 control trials (top) and 18 escapes out of 24 trials during optogenetic inhibition of VGAT^+^ neurons (bottom). Right: Single trial mouse tracking showing failed and successful escapes to shelter (red dots = escape termination) for control (top) and during iChloC stimulation (orange, bottom). **C**. Summary plot of escape probability in response to threat presentation with and without simultaneous optogenetic activation of VGAT^+^ dPAG neurons (n = 100 trials, N = 5 mice). **D**. Summary plot of escape probability in response to optogenetic inactivation and threat presentation (mean escape probability = 0.12 ± 0.09 Laser ON only, n = 108 trials, N = 4 mice). **E**. Left: Single mouse tracking showing control escape trials (Laser OFF) and trials with ChR2 stimulation *after* escape onset (red dot: escape termination; blue line: Laser ON). Right: Summary plots of the normalized mean escape termination distance from shelter during Laser OFF and Laser ON trials (N = 3 mice). Box-and-whisker plots show median, IQR and range. See also Figures S7 and S8.

## Discussion

We have described a population of GABAergic neurons in the dPAG that tonically fire action potentials and provide monosynaptic inhibition to dPAG excitatory neurons through activation of GABA_A_ receptors. In contrast with the sparse connectivity between VGluT2^+^ dPAG neurons (31), we find that these glutamatergic neurons receive input from local GABAergic dPAG neurons with a high connectivity rate, resulting in frequent inhibitory synaptic events that generate a sustained barrage of inhibition. The presence and role of a tonic inhibitory mechanism in the PAG was postulated decades ago (24) and recent work has described both longrange and local disinhibitory motifs as a main element of instinctive behaviour initiation (28, 42, 51, 52, 61–63). Our results add to the emerging view that sustained inhibition is a key property inherent to the functional architecture of the PAG and demonstrate its role in instinctive escape behaviour.

Our neural activity recordings from freely behaving mice show that VGAT^+^ dPAG are active as animals explore the environment, and that approximately half of the recorded population shows a transient decrease in activity before escape onset. We think that this could serve to disinhibit the VGluT2^+^ dPAG neurons that drive escape (31, 64, 65) and facilitate escape initiation. This view is strongly supported by the optogenetic silencing experiments, in which inhibition of VGAT^+^ dPAG neurons increased escape probability. Together with the finding that optogenetic activation of the same neuron population can decrease escape probability, these results show that VGAT^+^ dPAG neurons are a key node for determining the escape initiation threshold, in addition to the synaptic properties between the threat input circuits and the dPAG (31). The existence of a sustained source of inhibition in the dPAG might be important for dampening the excitability of the excitatory neurons and prevent initiation of escape behaviour in the absence of threat input, while also providing a means for fast, bidirectional modulation of escape. It will be interesting in the future to determine whether changes to the baseline firing of VGAT^+^ dPAG neurons are a mechanism to support plastic changes to the escape threshold, such as during learned suppression of escape (4). It will also be important to establish whether the function of VGAT^+^ dPAG neurons in escape is exclusively mediated by intra-PAG inhibition or whether they also directly modulate other targets, such as the cuneiform nucleus of the mesencephalic locomotor region, which has been previously reported to play a role in defensive behaviours (66–68).

In addition to setting a threshold for initiating escape behaviour, our results suggest that VGAT^+^ dPAG neurons are also important for the ongoing execution of escape, in particular for terminating the flight action. A large fraction of this population shows a gradual increase in activity between escape onset and escape termination at the shelter, and optogenetically increasing or decreasing their activity can cause animals to undershoot or overshoot their target, respectively. Given the known role of glutamatergic dPAG neurons in locomotor initiation and speed control during escape (31, 34, 48, 50, 65), these data can be interpreted as VGAT^+^ dPAG neurons acting to shut down firing in the excitatory neurons that drive escape. These results are in line with the previous finding that VGAT^+^ neurons in the lateral PAG have a role in both the initiation and execution phases of hunting behaviour (35). An important unanswered question is whether there are distinct subsets of VGAT^+^ dPAG neurons, one for controlling initiation and the other for execution. Our analysis identified two broad clusters based on the activity profile during escape, which may arise if there are groups of VGAT^+^ neurons integrating different input pathways (35). It will be interesting to determine if such input segregation exists. The observation that we were able to change both escape initiation and execution by manipulating activity of the entire population is consistent with the main output of the two clusters being glutamatergic dPAG neurons, with the timing of VGAT^+^ neuron activation being the main determinant of the effect on escape behaviour.

While the activity of glutamatergic dPAG neurons is positively correlated to escape speed (31), we found that the population activity of GABAergic neurons is not significantly modulated by speed but instead peaks at escape termination. This observation supports a general framework for locomotor-correlated activity in brainstem regions, where glutamatergic neurons are the key population for regulating speed and are under the control of local inhibitory networks (69). The finding that GABAergic cells in the dPAG mediate the termination of escape behaviour further adds to the view that the dPAG is an integral part of the supraspinal circuits for the coordination of goal-directed locomotion (69–72).

Studies of escape behaviour in the wild have highlighted how flexible the flight action must be since animals often engage in a complex series of stop-and-go events while computing and executing escape trajectories directed to safe locations (1, 10, 19, 73–75). Our results raise the hypothesis that inhibitory neurons in the dPAG might integrate spatial and sensory information to implement flexible speed adjustments during escape and determine when safety has been reached through modulation of their firing rates (41, 76, 77).

## Supporting information

Video_iChloc_threatOnset_Fig3

Video_ChR2_threatOnset_Fig3

Video_ChR2_controls_FigS7

Video_ChR2_beforeThreatOnset_FigS7

Video_ChR2_termination_Fig3

Video_iChloc_overshoot_FigS8

## Acknowledgements

This work was funded by a Wellcome Senior Research Fellowship (214352/Z/18/Z), by the Sainsbury Wellcome Centre Core Grant from the Gatsby Charitable Foundation and Wellcome (GAT3755 and 219627/Z/19/Z) and by a European Research Council grant (Consolidator no. 864912) (T.B.), German Research Foundation postdoctoral fellowships (E.K., project no. 515465001; A.V.S., project no. STE 2605/1), the UCL Wellcome 4-year PhD Programme in Neuroscience (O.P.A.), the SWC PhD Programme (F.C., S.C.L.) and the Max Planck Society (E.K., A.V.S.). We thank P. Iordanidou, T. Okbinoglu and M. Strom for technical assistance; the SWC Neurobiological Research Facility and MPI for Brain Research animal facility for animal care, FabLabs for technical support, and K. Betsios for programming the data acquisition software. We thank F. Rau, P. Zatka-Haas, A. Fratzl and all members of the Branco lab for discussions; D. Campagner, J. Kohl, S. Keshavarzi for comments on the manuscript. This manuscript was typset using a modified version of the HenriquesLab bioRxiv template.

## Author Contributions

A.V.S. and T.B. conceived the project with critical input from D.A.E. A.V.S, O.P.A. and E.K. performed *in vitro* electrophysiology. A.V.S. and D.A.E. performed *in vivo* calcium imaging and optogenetic manipulation experiments, F.C. contributed to preprocessing and analysis of the calcium imaging dataset. S.C.L. performed *in vivo* single-unit recordings, supervised by T.W.M. A.V.S and T.B. supervised the project. A.V.S., T.B. and D.A.E. wrote the manuscript and all authors read and commented on the manuscript.

## Competing Financial Interests

The authors declare there are no competing interests.

## Methods

### Animals

Adult (6–14 weeks old), male and female VGAT-ires-Cre (Jackson Laboratory, stock #028862), VGluT2-ires-Cre (Jackson Laboratory, stock #016963), crossed VGAT::tdTomato (Jackson Laboratory, stock #007908) and VGAT::eYFP or VGluT2::eYFP (Jackson Laboratory, stock #006148) mice were housed with ad libitum access to water and chow on a 12 h reversed light cycle and tested during the light phase. A subset of in vitro patch-clamp experiments was made from Gad2-T2a-NLSmCherry animals (Jackson Laboratory, stock #023140) (see below). Silicon probe recordings were performed in 8–10 weeks old, male C57BL/6J mice, obtained from commercial suppliers (Charles River). Animals were used for scientific purposes in accordance with the UK Animals (Scientific Procedures) Act of 1986 and the Animal Welfare and Ethical Review Body (AWERB), guidelines stated in Directive 2010/63/EU of the European Parliament on the protection of animals used for scientific purposes, institutional guidelines and following approval by the relevant authorities.

### General surgical procedures

Drugs were prepared on the day of surgery, diluted in sterile 0.9 % NaCl saline and warmed before use. Animals were anaesthetised with an intraperitoneal (i.p.) injection of ketamine (100 mg/kg) and xylazine (12 mg/kg), and carprofen (5 mg/kg) was administered subcutaneously for perioperative analgesic care. The hair on the forehead was shaved using a trimmer (Contura, Wella), and the skin disinfected. Isoflurane (0.5–1.5 % in oxygen, 1 L/min) was used to maintain anaesthesia. The skin was then cut to expose the skull, and a large craniotomy was made using a hand-held dental drill using a 0.5 or 0.7 mm stainless steel burr (Fine Science Tools) to expose an area between the confluence of transverse and sagittal sinus, and the anatomical border between the superior and inferior colliculi (AP from lambda: +0.1 to -1.0 mm; ML: ± 0.6 mm). A durotomy was performed using a 30G hypodermic needle. The connective tissue covering the confluence of sagittal and transverse sinus was gently pulled forward anteriorly using fine forceps to reveal the underlying superior colliculi. Viral vectors were delivered using a pulled cut glass pipette (10µl Wiretrol II with a Sutter P-1000) that was mounted on a stereotaxic injector (Janelia Research Campus, Ronal Tool Company) coupled to an oil hydraulic micromanipulator (MO-10, Narishige) on a stereotaxic frame (Model 963, Kopf Instruments), at ∼20 nl/min. After injection, the needle was kept at the site of injection for 10–20 min, before being retracted. Fibre optic and GRIN lens cannula implants were first slowly lowered to 50 µm beyond their final location, and then retracted. For lens cannula implants, a single 400 µm diameter fibre optic with a 60º conical tip was pre-inserted at the desired location and withdrawn, to allow for easier successive insertion of the 500 µm lens implant. The craniotomy was covered with a silicone sealant (Kwik-Cast, WPI) before implants were secured using light-cured dental cement (RelyX Unicem 2, 3M). The skin wound was glued (Vetbond, 3M) or sutured (6-0, Vicryl Rapide). To ensure maximal stability during calcium imaging, the fissures between the interparietal and occipital plates were glued after the implantation of GRIN lens cannulae and before wound closure. Coordinate measurements are given from lambda and skull surface. For precise rostrocaudal and mediolateral targeting of fibre placements and viral injections, coordinates were confirmed and, if necessary, re-aligned after the craniotomy and durotomy were performed based on the border between the superior and inferior colliculi (∼0.65–0.8 mm behind lambda) and the small midline blood vessel that runs sagittal between the two superior colliculi.

### Viruses

The following viruses were used in this study and are referred to by abbreviations in the text. For chemogenetic inhibition experiments, AAV2/2-hsyn-DIO-hM4D(Gi)-mCherry was obtained from Addgene (plasmid #44362, 4.6x10^12^ GC/ml). For optogenetic activation, AAV2/2-EF1a-DIO-hChR2(H134R)-EYFP-WPRE (3.9x10^12^ GC/ml) or AAV2/2-EF1a-DIO-hChR2(H134R)-mCherry-WPRE (6.6x10^12^ GC/ml; Deisseroth, UNC Gene Therapy Vectore Core) were used. Optogenetic inhibition experiments were performed with AAV2/9-EF1a-DIO-iChloC-2A-tDimer (3.75x10^12^ GC/ml; a gift from S. Wiegert and T. Oertner) or AAV2/1-EF1a-DIO-iChloC-2A-dsRed (5x10^13^ GC/ml; Addgene plasmid #70762, a gift from T. Margrie). For calcium imaging experiments, AAV2/1-hsyn-DIO-GCaMP6s-WPRE (6.25x10^12^ GC/ml) was acquired from Penn Vector Core.

### Electrophysiology recordings *in vitro*

Patch-clamp recordings were performed from identified glutamatergic and GABAergic neurons in acute, coronal slices of the midbrain containing the PAG and as previously described (31, 78).

#### Slice preparation

Slices were prepared by one of two alternative slicing methods (see lab protocol by Pavon Arócas and Branco (78) for details). In brief, animals were either anaesthetised using isoflurane and decapitated directly (path A) or after intracardial perfusion with ice-cold artificial cerebrospinal fluid (aCSF) (path B). Path A: the brain was extracted and submerged in carbogenated ice-cold sucrose-based aCSF containing (in mM): 87 NaCl, 26 NaHCO_3_, 50 sucrose, 10 glucose, 2.5 KCl, 1.25 NaH_2_PO_4_, 3 MgCl_2_, 0.5 CaCl_2_. Tissue blocks containing the midbrain PAG were mounted on a vibratome (VT1200S, Leica) and coronal slices (-4.8 to -4.3 mm from bregma) of 250–300 µm thickness were cut at 0.05–0.06 mm/s. After cutting, slices were kept in sucrose-based aCSF at near-physiological temperature (35º C) for 30 min before being transferred to a beaker with 35º C-warm aCSF (containing in mM: 125 NaCl, 26 NaHCO_3_, 10 glucose, 2.5 KCl, 2 CaCl_2_, 1 MgCl_2_, 1 NaH_2_PO_4_) that was then stored at room temperature for at least half an hour before recordings began. Path B: For a subset of experiments, the slicing and recovery aCSF contained (in mM): 96 NMDG, 96 HCl, 25 NaHCO_3_, 25 D-Glucose, 20 HEPES, 5 Sodium L-ascorbate, 12 N-actyl-L-cystein, 10 MgSO_4_, 3 Sodium pyrate, 3 Myo-inositol, 2 Thiourea, 1.25 NaH_2_PO_4_, 0.5 Ca_2_Cl, 0.01 Taurine. Slices were kept at 32–34º C to recover for a period of <15 min and then held in HEPESbased aCSF until recordings began. For both paths, slices were stored at RT for ∼4–5 hours after preparation before being discarded and all aCSF was equilibrated with 95% O_2_, 5% CO_2_.

#### General setup

Individual midbrain slices containing the PAG were transferred to a submerged chamber, perfused with ACSF heated to 32–33º C at a rate of 2–3 ml/min and visualised on an upright microscope with oblique contrast (Slicescope, Scientifica) using a 60x or 40x objective (Olympus). VGluT2- and VGAT-positive dPAG neurons were identified based on location, fluorescence protein expression and cell-attached firing properties. In a subset of recordings, putative glutamatergic neurons were identified by being VGAT- or GAD2-negative (not expressing a fluorophore; mCherry or eYFP) and non-spiking in cell-attached configuration before break-in. Given that all GABAergic neurons are tonically spiking and excitatory neurons are not, we consider this to be a reliable method to identify putative excitatory dPAG neurons. Patch-clamp recordings were performed with an EPC 800 (HEKA) amplifier and monitored using an HMO1002 oscilloscope (Rohde & Schwarz). Signals were filtered at 5 kHz, digitised with 16-bit resolution at 25 kHz using a PCI 6035E A/D board and recorded in LabVIEW (both National Instruments) using custom software. A subset of recordings was performed and digitised using a Multiclamp 700B amplifier with an Axon Digidata 1550B Digitiser, and recorded using Clampex (Molecular Devices).

#### Whole-cell and loose-seal patch-clamp recordings

Pipettes were pulled from borosilicate filamented glass capillaries (Harvard Apparatus, 1.5 mm OD, 0.85 mm ID) with a P-10 or PC-100 Narishige puller to a final resistance of ∼4–7 MO. Unless stated otherwise, for whole-cell patchclamp recordings, pipettes were backfilled with filtered internal solution containing (in mM): 130 KMeSO_3_, 10 KCl, 10 HEPES, 5 Na-phosphocreatine, 4 Mg-ATP, 1 EGTA, 0.5 Na_2_-GTP, 285–290mOsm, pH was adjusted to 7.35 with KOH. In some recordings, 1 mg/ml biocytin or neurobiotin was added on the day. To analyse inhibitory postsynaptic currents (IPSCs) onto VGluT2 neurons, a high chloride internal solution was used containing in mM: 155 KCl, 10 HEPES, 2 EGTA, 2 Na-ATP, 2 Mg-ATP, 0.3 Na-GTP or 140 KCl, 10 HEPES, 5 Na-phosphocreatine, 2 Mg-ATP, 0.5 Na-GTP, 0.2 EGTA, 285–290mOsm, pH adjusted to 7.35 with KOH. For loose-seal cell-attached recordings, pipettes were backfilled with aCSF and a seal with a resistance of 10–20 MO was established. Action potential currents in voltage-clamp mode were recorded continuously for 5–6 min. A holding potential of 0 mV was maintained through current injections (within a range of ± 20 pA) and the access resistance was monitored throughout the recording. For whole-cell recordings, the resting membrane potential was determined in current clamp directly after establishing the whole-cell configuration and experiments were only continued if cells had a resting membrane potential more hyperpolarised than -45 mV (not corrected for liquid junction potential). In a subset of recordings, cells were further characterised by recording their membrane responses to current injections of increasing strengths (-40 to + 140 pA, increment: 20 pA, 500 ms duration). The access resistance was monitored continuously, and was compensated in current-clamp recordings. Only cells with a stable access resistance < 30 MO were analysed. Voltage-clamp recordings were performed at a holding potential of -60 mV with the exception of IPSC recordings using the high-chloride internal solution, which were performed at -70 mV.

#### Stimulation

Electrical stimulation was performed using a stimulus isolator (DS2A, Digitimer) with a pipette placed in the deeper layers of the superior colliculus. Pulse trains were delivered every 15 s with a stimulation intensity of 0.02–4.3 mA, a pulse duration of 100 µs and an interpulse interval of 50 ms. For optogenetic stimulation experiments, recordings were made > 30 days after injection of AAV2-DIO-ChR2mCherry into the dPAG of VGAT-Cre mice. Wide-field 490 nm LED illumination (pe-100, CoolLED) was used to excite ChR2 expressed in VGAT dPAG neurons. Pulse trains (5 pulses, 50 ms interpulse interval, 1 ms pulse duration, maximum light intensity = 2.7 mW) were delivered every 30 s. The stimulation intensity for both electrical and optogenetic stimulation was adjusted for each experiment.

#### Pharmacology

To block glutamatergic synaptic transmission, kynurenic acid (Sigma) was added to the aCSF as powder to a final concentration of 2 mM and sonicated until dissolved. GABA_A_ and glycinergic receptor-mediated synaptic transmission was blocked using 50 µM picrotoxin (Tocris Bioscience), which was prepared from a 100 mM stock solution containing DMSO (final DMSO concentration: 1/2000). GABA_A_ receptor-mediated synaptic transmission was blocked using 10 µM SR-95531 hydrobromide (Gabazine, Hellobio). Water-soluble clozapine N-oxide dihydrochloride (CNO, Hellobio) was bath applied at a final concentration of 10 µM from a 100 mM stock solution. Control recordings (matched in duration to experiments with bath application of drugs) were performed interleaved and, if possible, in the same animals. Drugs were added to the aCSF as stated in the text or figure legends.

#### Data analysis

Membrane potentials were not corrected for liquid junction potential. The spontaneous firing frequency of neurons recorded in loose-seal cell-attached configuration was analysed in a period of 2–4 min after seal formation. For whole-cell recordings, the resting membrane potential was calculated during current steps in a 100 ms baseline window in current clamp mode. The input resistance (R_in_) was calculated from the steady-state voltage measured in response to a hyperpolarising test pulse (-40 pA, 500 ms duration) at holding potential of -60 mV. The membrane time constant was calculated by fitting a single exponential to the decay of the test pulse (y = y0 + Ae^(-(x-x0)/τ)^). The mean action potential firing frequency of VGAT neurons in response to current injection of increasing strengths (20–140 pA, 20 pA increments) was calculated from two consecutive repetitions per cell. Spontaneous inhibitory postsynaptic currents (IPSCs) onto glutamatergic dPAG neurons were detected using a threshold algorithm, and their frequency and peak amplitude were analysed from a continuous 30 s time window every 1 min or continuously and binned by minute. The peak phasic conductance was calculated from the cell’s mean peak current amplitude divided by the holding potential (G = I / V_hold_). The mean unitary charge was calculated from the average spontaneous IPSC for each cell, integrating the area over a 50 ms window from IPSC onset. The average charge was estimated by multiplying the unitary charge by the frequency of events for that cell. For estimating the change in IPSC frequency in SR-95531 and CNO respectively, drugs were bath applied after a stable baseline recording of 2–5 min and recordings were continued for > 5 min. The mean baseline IPSC frequency was calculated from 2 min before bath application of the drug, and the mean IPSC frequency after drug wash-in was calculated for min 7–8 for CNO and min 5–7 for SR-95531 and their respective controls. Pharmacological effects of each drug were statistically assessed in comparison to time-matched controls (see above). Mean peak amplitudes of evoked postsynaptic currents were calculated from averages of a minimum of 5 sweeps both for baseline and drug conditions (where applicable). The paired pulse ratio of a synaptic input was calculated by dividing the mean peak amplitude of the second pulse by the first pulse. Statistical tests were performed on all cells pooled across animals. The coefficient of variation was calculated for each cell as the standard deviation of the inter-spike intervals (ISI) over the mean ISI. CV_2_ was calculated after Holt et al. (55) as:

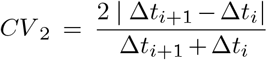

for each spike *i*, and the average CV_2_ was taken for each cell

### Behavioural recordings

#### Setup and acquisition

Behavioural experiments were performed in a rectangular arena made from Perspex (L: 60–80 cm × W: 20 cm × H: 40cm) with an infrared light-transmitting red shelter (L: 10 cm × W: 19 cm × H; 13.5 cm) in one corner of the arena, and as previously described (31). Briefly, a screen made of 100 µm thick drafting film (90 cm × 70 cm, Elmstock) was suspended 64 cm above the arena floor, and an ultra-short throw projector (PF1000U, LG) was used to project a grey background onto the screen, providing a uniform luminance of 3–4 lux (measured on the floor of the arena). Behaviour was recorded at 50 frames per second with a near-infrared camera (acA1300-60gmNIR GigE, Basler) positioned above the arena centre, flush with the screen. Video recordings, as well as sensory and optogenetic stimulation protocols, were controlled with custom-written software (mantis64) in LabVIEW (2015 64-bit, National Instruments). A PCIe-6351 board (National Instruments) was used to trigger and synchronise all signals. Sensory stimuli were delivered when the mouse entered a predefined ‘threat area’ at the opposite end of the arena to the shelter (∼15–20 × 20 cm). A 35 mm Petri dish was positioned in the threat area to enrich the environment and encourage exploration. Animals were allowed to explore the arena for a minimum of 7 min at the beginning of the recording, and a behavioural session typically lasted between 30 min and 2 hours.

#### Sensory stimulation

Escape was elicited using both auditory and visual stimuli. A frequency-modulated upsweep from 17 to 20 kHz over 3 s was used as an auditory stimulus, and delivered at different intensities ranging from 60 to 90 dB above the threat area via an ultrasound speaker (L60, Pettersson). The visual stimulus consisted of a sequence of 3 or 5 dark, linearly expanding circles (visual angle min– max: 2.6–47º, linear expansion: 118º / s over 260 ms, size maintained for 260 ms at the maximum visual angle) with an inter-spot interval of 500 ms. The contrast of the spots ranged between a Weber contrast of 60, 75 and 98 % (from low to high, converted to percentage from the respective negative Weber fractions), and was adjusted by changing the intensity of the circle against the grey screen which was maintained at a constant luminosity (see above). Visual stimuli were predominantly used and interleaved with auditory stimuli to prevent habituation. The stimulus intensity was adjusted based on the escape probability of each mouse in each session and the experimental paradigm (as described in the main text).

#### Behavioural data analysis

Mouse coordinates were extracted from the behavioural videos using DeepLabCut (79). Behavioural events were scored and annotated manually, including for escape: (1) first stimulus-reaction response after sensory stimulation (usually an ear movement followed by a brief startle), (2) beginning of the shelter-directed head-turn followed by a full-body turn, (3) start of run towards shelter, (4) escape termination. A successful escape was defined as a sequence of events (1) – (4), with an *uninterrupted* run into the shelter within 6 s of the end of a sensory stimulus (7.5 s for freely-moving calcium recordings to account for head-mounted microscope weight). The escape probability for a given stimulus is the fraction of trials which led to an escape to the shelter and was calculated for each animal separately. The peak velocity of the escape was determined by analysing the speed trace from the initiation of the escape until entry into the shelter.

### Optogenetic experiments in freely moving animals

#### Surgeries

For optogenetic activation experiments, VGAT-Cre mice were injected with AAV-DIO-ChR2 into the dPAG (1–3 injections per hemisphere with a combined volume of 40–150nl per side; ML: ±0.4 to 0.45, AP: 4.6 to -4.9, DV: -2.2). Dual optic fibres with 60º conical tips (400 µm diameter, 1.2 mm apart, DFC_400/430-0.48_3.5mm_GS1.2_C60; Doric Lenses Inc.) were implanted above the centre of the injection sites (AP: -4.6 to -4.9, DV: -1.9 to -2.0). Behavioural testing was performed 13–65 days after surgery (mean = 31.1 ± 4.3 days), with 2– 5 sessions per animal. For optogenetic inactivation experiments, VGAT-Cre mice were injected with AAV-DIO-iChloC into the dPAG (1–2 injections per hemisphere with a combined volume of 100–180nl per side; ML: ±0.4 to 0.45, AP: -4.6 to -4.9, DV: -2.2). Dual optic fibres with 400 µm thickness were implanted as described above (AP: -4.6 to -4.9, DV: -1.9). In two experiments, a single optic fibre (200-µm diameter, MFC-SMR; Doric Lenses Inc.) was implanted medially (ML: 0.0, AP: -4.6 to -4.9, DV: -1.9).

#### Behavioural testing

Behavioural testing was performed 12–40 days after surgery (mean = 23.8 ± 3.8 days), with 1–5 sessions per animal. Dual or single fibre-optic magnetic patch cords were attached to a fibre-optic rotary joint (FRJ_1×2i_FC-2FC_0.22 or FRJ1×1, both Doric Lenses Inc.) to allow unhindered movement. Light was delivered by a 472 nm laser module (Stradus, Vortran) with direct analogue modulation of laser power. Mice were presented with visual or auditory stimuli that elicited escape, and laser-on trials were interleaved with laser-off trials. Laser-on movement controls were performed without sensory stimulation. ChR2 stimulation was delivered at a frequency of 50 Hz for 3 or 5 s (pulse duration: 5 ms, laser intensity: 18.5–30 mW per hemisphere/optic fibre). Continuous blue-light stimulation of iChloC was delivered for 3 or 5 s at laser intensities of 0.03–0.3 mW using a neutral density filter (optical density: 0.3, Thorlabs) fitted in front of the laser output. Laser intensities were calibrated during exploratory movement to ensure no escape-like movements were elicited with too high laser power. To test for a bidirectional behavioural effect of VGAT neuron activation and inhibition, two separate behavioural paradigms were used to adjust the initial escape probability by changing the intensity of the visual (contrast) and auditory (dB) stimuli. Since we predicted that escape probability would decrease with VGAT neuron activation, we adjusted the sensory stimuli for each mouse during this experimental set to yield a high escape probability of ∼ 80%. Conversely, predicting an increase of escape probability with VGAT neuron inactivation, we used low intensity stimuli to achieve a baseline escape probability of ∼ 20–40%. To test the effect of optogenetic activation of VGAT neurons on escape termination, the laser was triggered manually immediately after the onset of stimulus-evoked escape runs. To test whether inactivation of VGAT neurons causes an increase in the escape distance, we moved the shelter away from the end wall into the middle of the arena. The shelter is open front and back to allow animals to pass through. The animals were allowed to explore the new shelter location, before a sensory stimulus was delivered to elicit escape. If the escape was to the new shelter location, indicating that the animal had memorized the new shelter location, we proceeded to interleave the sensory stimulation with concomitant iChloC activation trials.

## Data analysis

Behavioural tracking data was annotated and analysed as described above. The effect of optogenetic activation of VGAT^+^ neurons on escape termination is calculated as the normalized distance from the shelter (0 = shelter entrance, 1 = far end of arena) at which the animal stopped its escape run. An average of all trials per tested animal is reported.

### Calcium imaging in freely moving animals

#### Surgeries and acquisition

Calcium imaging experiments were performed as previously described (31). Briefly, a miniaturised head-mounted fluorescence microscope (Model L, Doric Lenses Inc., Canada) was used to image the genetically encoded calcium sensor GCaMP6s in VGAT neurons of the dPAG. A volume of 100 nl AAV2/1hsyn-DIO-GCaMP6s was injected into the dPAG (AP: -0.45 to -0.7, ML: +0.25–0.3, DV: -2.2). A 500 µm diameter GRIN lens imaging cannula (Model L, Doric Lenses Inc.) with an 80 µm working distance was slowly inserted, with the final tip position ∼100 µm above the centre of the viral injection site. At least 19 days after surgery, the microscope was attached to the snap-in cannula implant on the mouse’s head without anaesthesia and the mouse placed in the behavioural arena. Behavioural recordings were performed only after ensuring a clear field-of-view (FOV) was present (mean first session = 35.1 ± 5.3 days, mean last session = 62.2 ± 9.3 days after surgery; range = 19–104 days, N = 8 mice). Data were acquired with a resolution of 630 *×* 630 pixels at 10–20 Hz, with an excitation power of 150–300 µW and with a gain of 1 or 2. Sensory stimulation was delivered as described above.

#### Data analysis

Fluorescence movies were downsampled from 630 *×* 630 to 126 *×* 126 pixels and then flatfield corrected. To obtain an estimate of the movie’s flatfield, we calculated the average intensity across time and applied a gaussian blur to the resulting image. The flat-field was centred around zero by subtracting its mean and this image was then subtracted from all frames of the movie. The movies were cropped to remove vignetting at the edges of the recorded FOV. The open-source analysis pipeline CaImAn (80) was used for motion correction and component extraction and curation. Briefly, movies were motion corrected using the NoRMCorre algorithm that corrects non-rigid motion artifacts, and source extraction was performed using constrained non-negative matrix factorization to extract components with overlapping spatial footprints. The extracted components were manually curated (merged, rejected or added) after visual inspection and the components’ spatial shapes saved as cell masks. These masks were used to extract the background-subtracted (rolling-ball, 10px radius) raw fluorescence signal over time from the motion-corrected videos, and linearly interpolated to 30 Hz to match the behaviour camera frame-rate. Imaging sessions were almost always intermittent (mean 16 days between sessions with 1–3 sessions per 8 animals, range 1–72 days, one animal with three consecutive days), and precluding FOV registration between imaging sessions and repeated cell identification. Each FOV was therefore treated as an independent sample. Delta F/F and Z-scored activity was used as the basis of all further analysis. Traces with a Z-score above 1.96 at any time during the recording session were considered active neurons.

To analyse VGAT^+^ neuron activity in the freely-moving animal in the absence of threat, single neuron activity traces acquired during the baseline period of each session (before the first threat stimulus, mean duration 548 s, range 339–1046 s, n = 209 neurons, from 17 sessions in 8 animals) were Z-scored and Savitzky-Golay filtered (2 s window, polyorder = 2, *scipy*). Events were then detected using the *scipy*.*find_peaks* function (prominence = 0.3) to calculate the mean event rate per mouse. Neurons were further sorted by whether they were active during escape or not (see methods below), and whether events were coincident with animal movement, defined as the centre-of-mass of the animal exceeding 2 cm/s. For quantifying locomotion, movement periods were defined by timepoints when the animal’s speed was greater than 1.5 cm/s and stationary periods as less than 1.5 cm/s while the animal was outside the shelter in the arena, in the baseline period before the first threat stimulus. A locomotion modulation index (LMI) and significance tests were calculated after Henschke et al (57). The LMI was defined as the difference between the mean F/F_0_ during locomotion (R_L_) and stationary (R_s_) periods, normalized by the sum, for each neuron as LMI = (R_L_ – R_s_)/(R_L_ + R_s_). To determine significance for the modulation indices, bootstrapping with replacement was used to estimate the error for each neuron. The data was binned into 1 s periods of continuous locomotion or stationary periods. The mean ΔF/F_0_ was taken for each bin and regarded as a single sample. The average correlation between consecutive samples was found to be R = 0.57. We then randomly selected a reduced number of samples (43% = 1-R, to reflect the fact that samples are not completely independent) of ΔF/F_0_ with replacement from the original set of samples, repeating this 1000 times to obtain 95% confidence intervals for individual significance tests for each neuron. If a neuron’s 95% confidence interval was significantly different from an LMI of 0 and its absolute LMI value exceeded 0.2, indicating >50% change in F/F_0_ between locomotion and stationary states, it was considered significantly locomotion responsive.

To identify neurons active during escape trials, Z-scored activity traces were first baselined by subtracting the mean value of 500 ms preceding stimulus onset. For each neuron, the distribution of activity slopes for each trial event (linear regression to a 660 ms window of activity starting at the following event onsets: *stimulus, first stimulus-reaction response, escape turn, escape run, shelter entry, escape stop*) over all trials in a session, was compared to the distribution of activity slopes in the 660 ms baseline periods preceding stimulus onset (two-tailed t-test with Bonferroni correction, P = 0.05) to determine whether there was a statistically significant change in signal for any event in the stimulus-escape sequence for that neuron, followed by a manual inspection and curation. 39 neurons showed a motion artefact (sharp symmetrical downward deflection) as the microscope cables contacted the roof of the shelter during escape running, which was removed by cubic interpolation 500 ms either side of the artefact trough. To compare neural activity across all escape-to-shelter trials and all animals, where the sequence of behavioural events is stereotyped but inter-event intervals are variable, the median inter-event intervals for each behavioural event relative to stimulus onset was calculated and the median of these distributions was used to construct a template with a 2 s window added after escape-end. Z-scored calcium traces were then linearly interpolated to the template and the mean calcium trace calculated for each escape-active neuron (n = 123) for escape trials and fail-to-escape trials. As experiments were acquired at 10–20 Hz, all traces were identically low-pass FFT filtered to remove high-frequencies that were present in a subset of sessions. To cluster neurons by their activity profile, the mean calcium over time traces of all neurons (escape trails only) were first ternarized by double thresholding of activity (y < -0.2 Z = -1, y > 0.2 Z = 1, otherwise y = 0), transformed by principal component analysis, then K-means clustering was performed on the first 3 principal components (cumulatively explaining 86% of the dataset before thresholding into ternary traces) using n = 2 clusters, indicated as optimal by silhouette scores and elbow method. The timing of calcium activity relative to escape end was calculated using the time of peak or trough activity over the time warped mean trace for each neuron.

### Histological quantification of virus expression and fibre location

For *in vitro* electrophysiological recordings, the expression of virus was visually confirmed for each slice using 490 nm or 568 nm LED illumination (for green and red fluorescent proteins, respectively) prior to experimentation on the upright microscope. Slices from animals with clear infection of the deeper layers of the superior colliculus were discarded. For histological confirmation of fibre placement and injections site of *in vivo* calcium imaging and optogenetic experiments, mice were anaesthetized with pentobarbital (400–800 mg/kg, final volume: 0.1 ml per 10 g, diluted 1:1 in sterile 0.9 % NaCl solution) and decapitated. To remove optic fibres and GRIN lenses without damage to the brain, the dorsal skull was cut in a circle around the cemented fibres and was carefully removed together with the fibres. Brains were then extracted from the base of the skull and fixed in 0.1 M PBS containing 4% PFA overnight at 4º C. Slices of 100 µm thickness were cut in 0.1 M PBS on a HM650V vibratome (Microm), mounted using SlowFade Gold Antifade Mountant with DAPI (ThermoFisher) and imaged on a wide-field fluorescence microscope (AxioImager, Zeiss). The placement of optic fibres and GRIN lenses was assessed based on their tract and tip location. Tip locations are illustrated in the respective sections of the Paxinos mouse brain atlas (81) (see Figures S4 and S7). The expression of viruses was visually confirmed. For AAV2/2-DIO-ChR2 injections, only animals with expression localised to the dPAG were used. Animals with expression in the deeper layers of the SC were excluded. For AAV-DIO-iChloC injections (AAV2/1 or AAV2/9), experiments were included based on fibre tip location in the dPAG. Animals with tip locations outside the caudal dPAG (< -4.6 mm or > -5 mm from bregma) or tip locations not aligned with the virus injection were excluded.

### Chronic silicon probe recordings *in vivo*

#### Surgery

Mice were anaesthetised in an isofluorane induction chamber (3–4 % isofluorane, 1 L/min) before being transferred to a nose cone (2 % isofluorane, 1 L/min). Surgical procedures were performed under anaesthesia using isofluorane (1.5–2 %, 1 L/min). Protective gel was applied to the eyes (Puralube Vet Ointment), analgesia was given subcutaneously (Metacam 25 µl/10g) and the temperature was maintained at 36º Celsius using a heating pad and temperature probe. The mouse was then secured on a stereotaxic frame (Angle Two, Leica Biosystems. After incision, the skull was roughened and a small craniotomy was made away from the site of implantation and a ground pin inserted. Next a ∼1 mm diameter craniotomy was made behind lambda using a dental drill (Osada Electric, Japan) to expose the transverse sinus. Connective tissue and dura were removed and the vessel pushed forwards. A silicon probe (4-shank Neuropixels 2.0, IMEC)(82), coated with DiI (Invitrogen), was slowly inserted at a 30º angle, ∼1 µm / s for 3 mm using a micromanipulator (Luigs and Neumann). Once in place, the probe was attached to the skull using UV-curing dental cement (RelyX Unicem 2 Automix, 3M), reinforced with dental cement (Super Bond C&B, Sun Medical). The ground was then connected to the pin and the implant reinforced with dental cement. A protective casing was attached and cemented in place using UV-curing cement. Following surgery, the mouse was returned to its home cage to recover on a heat pad. Following recovery, the mouse was briefly anaesthetised and the headstage connected and fixed in place.

#### Data acquisition and visual stimulation

We used the same behavioural arena and testing protocols as described previously (4). Briefly, each mouse was transferred in their home cage to the experimental room and given at least 5 min to acclimatise to the room under low light conditions, before being temporarily anesthetised to connect the Neuropixels interface cable to the headstage. A rotary joint was used to allow cable de-rotation during recording (AHRJOE_1x1_PT FC_24, Doric Lenses). Mice were allowed to explore their home cage to acclimatise to the attachment and recover from anaesthesia, before being transferred to the behavioural arena (red Perspex with a white opaque floor, L:50 cm × W:20 cm × H:28 cm) with a red dome (10 cm diameter) or rectangular shelter (10 cm x 20 cm x 10 cm, both red Perspex). For chronic silicon probe recordings, data were acquired at 30 kHz using Neuropixels probes connected to a PXIe-1000 card (IMEC) housed in a PCIe-1071 chassis together with a PXIe-8398 remote control module connected to a PCIe-8398 host interface card (National Instruments). Data were acquired using SpikeGLX software. The 30 Hz TTL signal to trigger camera frame acquisition was recorded on the probe synchronisation channel.

#### Data pre-processing and analysis

Electrophysiological data were pre-processed using SpikeInterface (83) to implement phase shift correction, global common median referencing and bandpass filtering (300–6000Hz) and run spike sorting using Kilosort 2.5 (82, 84). Units were manually curated using Phy (https://github.com/cortex-lab/phy) to exclude noise and multiunit activity from further analysis. Each unit included in the analysis passed manual curation, had a signal to noise ratio greater than 2, an inter-spike interval violations ratio less than 0.15 and an amplitude cut-off proportion less than 0.1. Baseline spontaneous firing rates were measured from a 180 second window during free exploration of the behavioural arena.

#### Histological quantification of probe tracks

To determine the exact location of probe tracks and single unit positions in the brain, a 3D whole-brain dataset was obtained at the end of each experiment using serial section microscopy. Mice were anaesthetised with pentobarbital (final volume: 100–150 µl, diluted 1:3 in saline), transcardially perfused with 10–20 ml of 0.1 M PBS followed by 10 ml of 4% PFA (diluted in PBS). Brains were retrieved and fixed overnight in 4 % PFA before being transferred to PBS. Brains were then mounted in 5% agar, glued to a microscope slide and transferred to a serial section two-photon microscope (85, 86). The microscope was controlled using ScanImage (v5.6, Vidrio Technologies) with BakingTray, a custom software wrapper for setting imaging parameters (github.com/SainsburyWellcomeCentre/BakingTray, doi.org/10.5281/zenodo.3631609). The agar block was submerged in 50 mM PBS and the brain was imaged using a 920nm laser (Mai Tai eHP DS, Spectra-Physics). Dichroic mirrors and bandpass filters were used to separate red, green and blue (background channel) signals, detected using three PMT channels (R10699, Hamamatsu Photonics with DHPCA-100 amplifiers, Femto Messtechnik). The entire brain was imaged with 2–4 optical sections per 50 µm physical section. Images were assembled following acquisition using StitchIt (github.com/SainsburyWellcomeCentre/StitchIt, zenodo.org/badge/latestdoi/57851444) and brains were then registered to a standardised atlas containing the individual columns of the PAG (87). DiI tracks were segmented using Brainreg-Segment (88, 89) and the depth of units along the probe used to determine their coordinates in atlas space. Only units located in the dPAG were included in the analysis.

### General data analysis

Data analysis was performed using Python 3.6 and 2.7 using custom-written routines, unless stated otherwise. Code will be made available upon request. Data are reported as mean ± SEM unless indicated otherwise. Statistical comparisons were made as stated in the text. Data was tested for normality, and parametric or non-parametric tests were chosen accordingly and as stated in the text. Statistical tests were made in SciPy Stats and Prism 9 (GraphPad). In all cases, p < 0.05 was the statistical threshold. No statistical methods were used to predetermine sample sizes, and are similar to those reported in previous publications (31, 54). The experimenters were not blinded to the conditions of the experiment during data acquisition and analysis.

## Supplementary Figures

**Figure S1.**
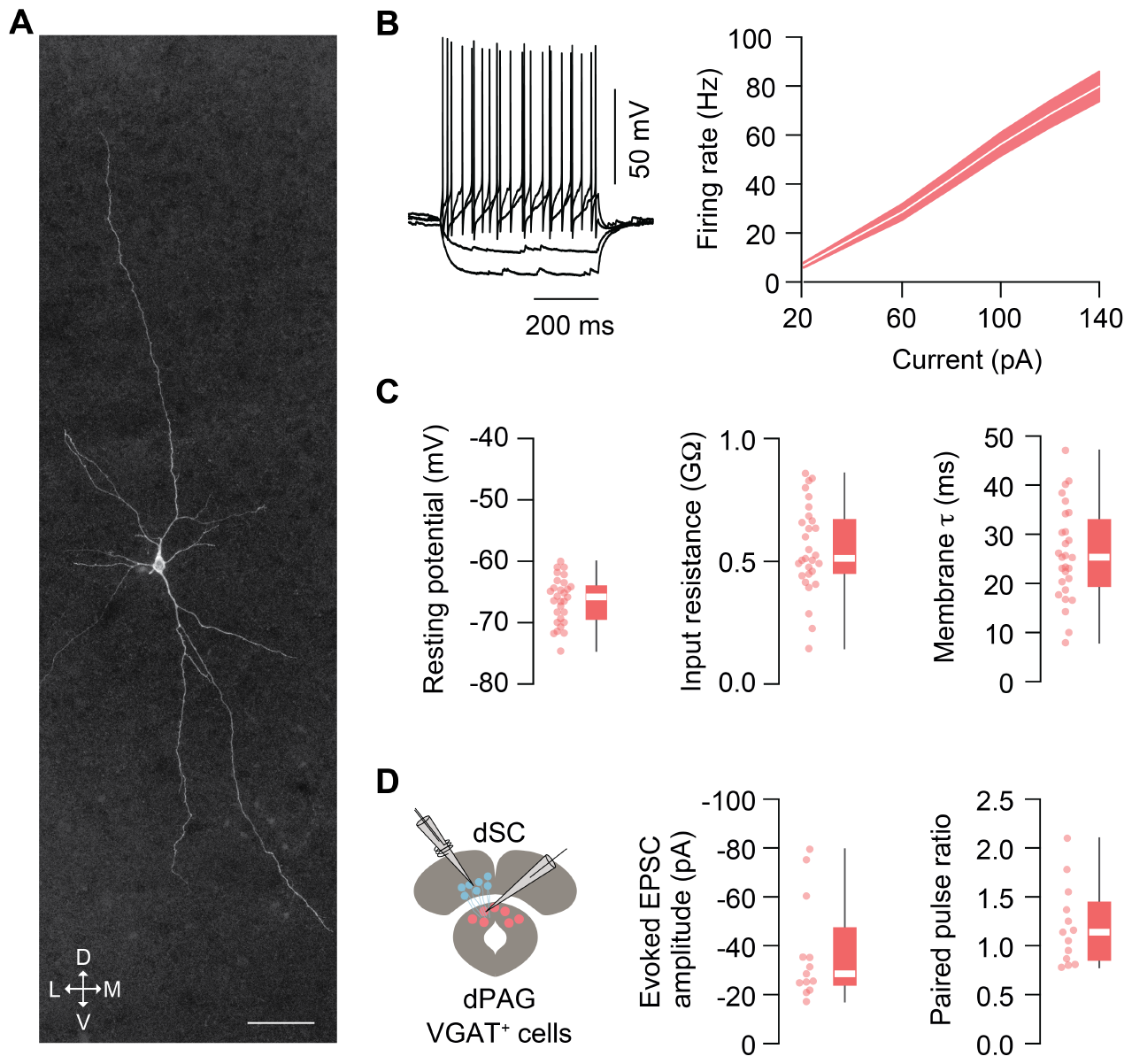
Electrophysiological properties of VGAT^+^ dPAG neurons. **A**. Maximum intensity projection of a confocal stack of a biocytin-filled VGAT^+^ neuron in the dPAG. Arrows indicate the neuron’s orientation: D – dorsal, V – ventral, M – medial, L – lateral; scale bar = 50 µm. **B**. Left: membrane potential traces of a VGAT^+^ dPAG neuron in response to current injections of different amplitudes. Right: summary graph of the current-firing rate relationship of VGAT^+^ dPAG neurons (shaded area is s.e.m; n = 29 cells, N = 8 mice). **C**. The mean resting membrane potential (-66.31 ± 0.69 mV), input resistance (0.55 ± 0.033 GΩ) and membrane time constant (25.61 ± 1.79 ms) were calculated for the same neurons as in B. **D**. Electrical stimulation of the deeper layers of the superior colliculus (dSC; left panel: schematic of the experimental configuration) evokes excitatory postsynaptic currents in all tested VGAT^+^ dPAG neurons with a mean peak amplitude of -37.07 ± 5.78 pA (peak amplitude of first pulse; middle panel) and a mean paired pulse ratio of 1.2 ± 0.11 (right panel) (n = 13 cells, N = 4 mice). Box-and-whisker plots show median, IQR and range, as well as individual data points.

**Figure S2.**
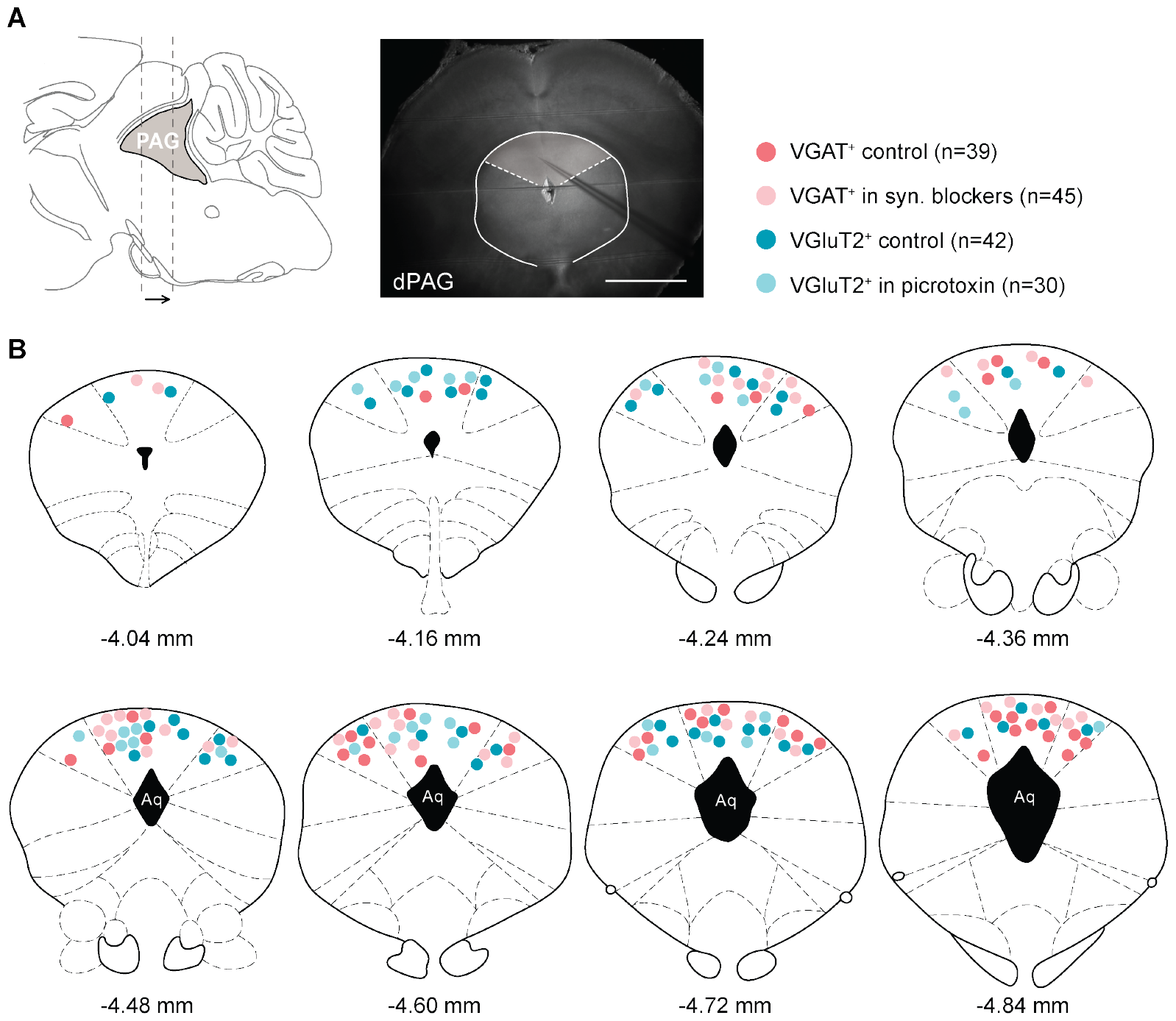
Anatomical location of loose cell-attached recorded cells in acute coronal slices of the dPAG. **A**. Schematic of a sagittal brain section indicating the rostrocaudal extend of recorded neurons in the PAG (left panel), and an image of an acute coronal midbrain slice with the pipette at its recording location in the dPAG (scale bar = 1 mm; right panel). Recordings were made from identified VGAT^+^ (filled circles, dark and light red) and VGluT2^+^ neurons (filled circles, dark and light blue; see legend). **B**. Superimposition of individual recorded cells along the rostrocaudal axis of the PAG, coordinates are in mm and from bregma. Mouse brain images adapted from (Paxinos and Franklin, 2001).

**Figure S3.**
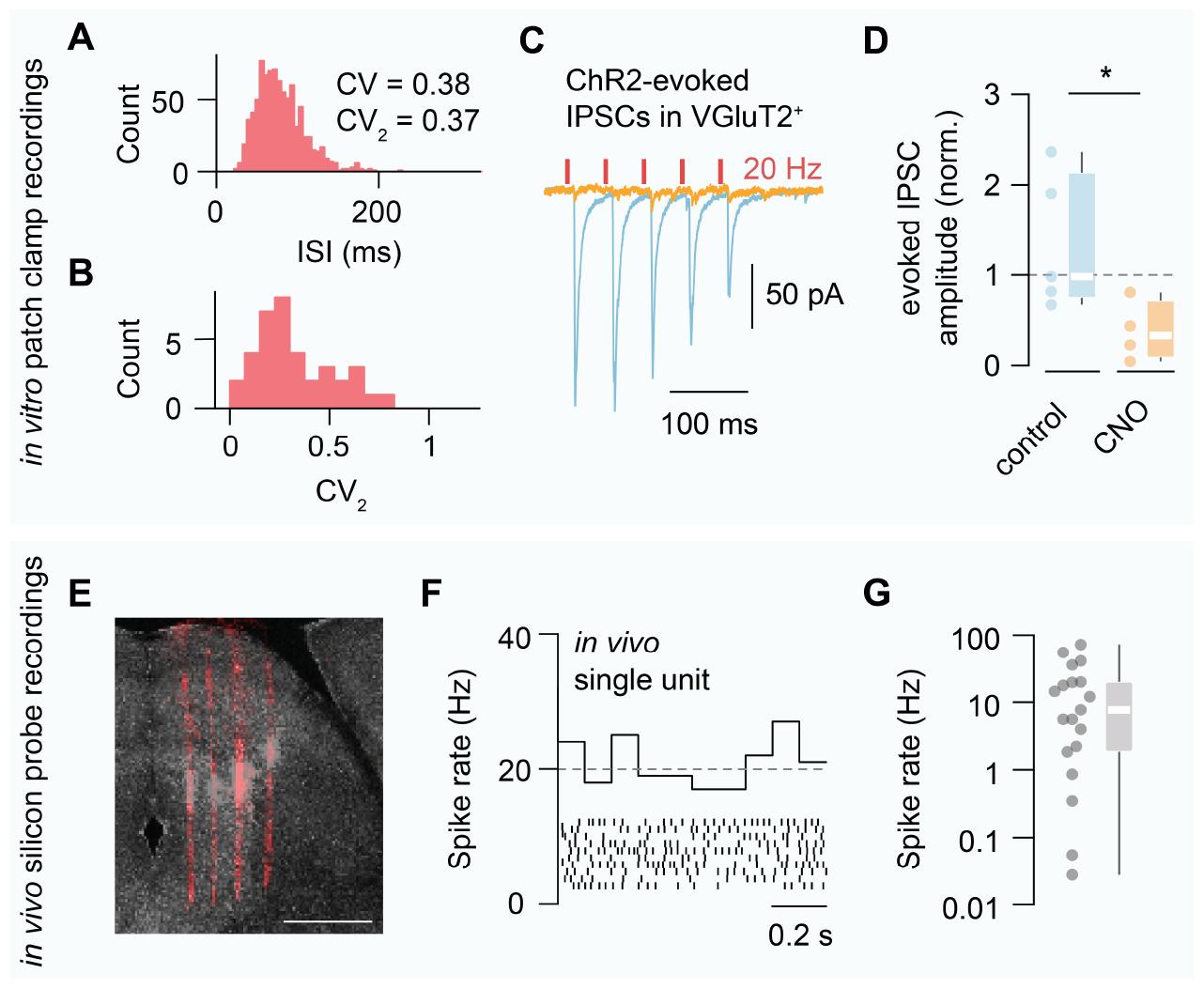
Firing properties of VGAT^+^ dPAG neurons and IPSCs *in vitro* and firing rates *in vivo*. **A**. Example ISI histogram of a regularly firing VGAT^+^ neuron recorded *in vitro*. **B**. Summary histogram of the mean CV_2_ of all VGAT^+^ dPAG cells recorded in control conditions. **C**. Example trace of ChR2-evoked IPSCs recorded in a putative dPAG excitatory neuron (stimulation protocol: 1ms duration, 5 pulses, 20Hz) before (blue trace) and after application of 10 µM CNO (orange trace) to silence VGAT^+^ dPAG neurons. **D**. Summary plot of the normalized evoked IPSC amplitude in control conditions and after 10 µM CNO application (control: n = 5 cells; 10 µM CNO: n = 4 cells; N = 2 mice). **E**. Example image of a coronal midbrain slice with the tracks of a 4-shank Neuropixels 2.0 probe made visible through DiI staining of the shanks prior to insertion into the brain (scale bar = 1 mm) . **F**. Example firing of a single unit recorded in the dPAG *in vivo* during exploration with the mean spike rate (histogram) over time shown for 10 randomly sampled 1 s time intervals from within a 60 s time window. The grey dotted line indicates the mean spike rate (19.96 Hz) for the same neuron. **G**. Average spike rate of all dPAG single units recorded *in vivo* (note log scale of y-axis). Each plotted data point (grey filled circles) is the mean firing rate of a single unit. Box-and-whisker plots show median, IQR and range, as well as individual data points.

**Figure S4.**
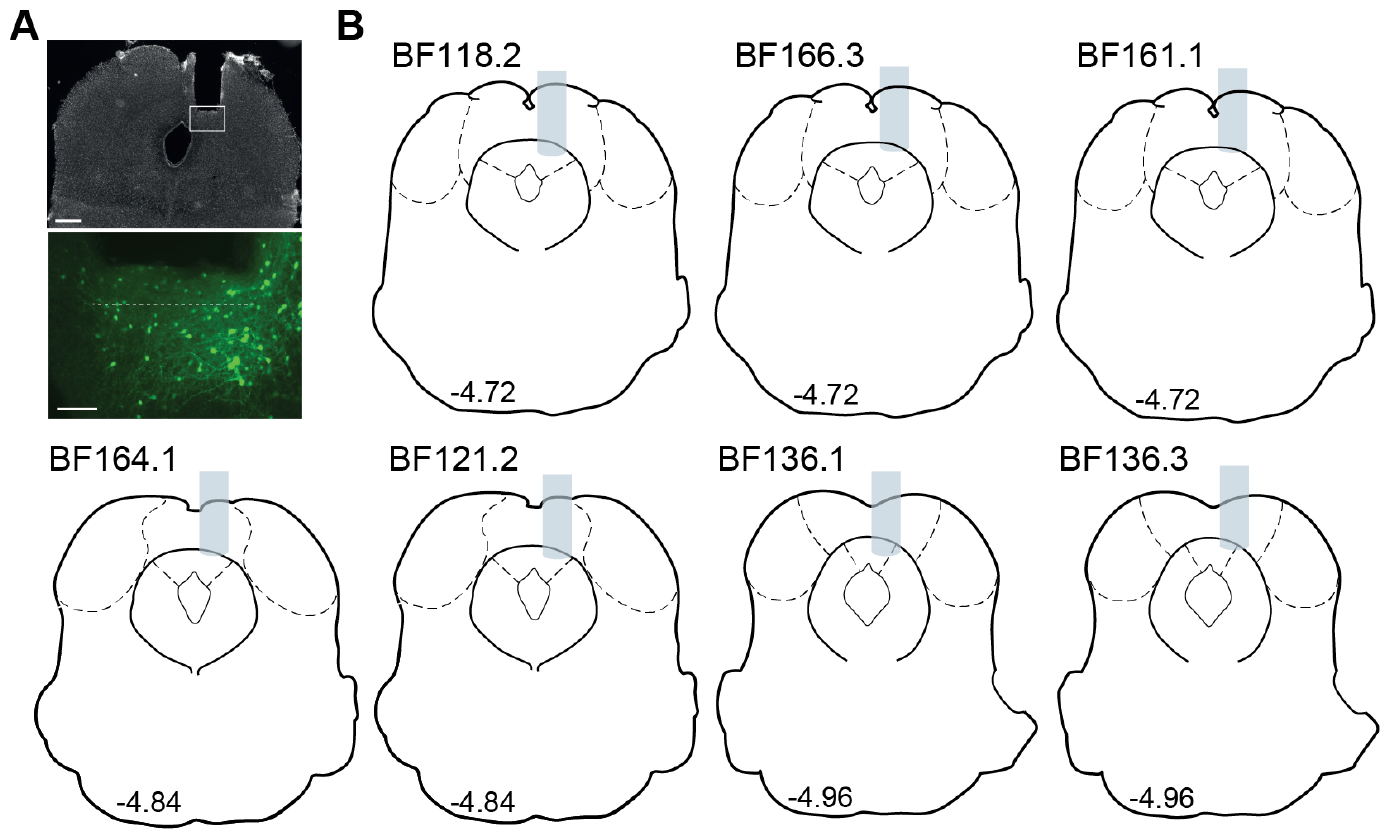
Gradient index lens placements in the dPAG for *in vivo* calcium imaging recordings using a miniaturised microscope in freely moving mice. **A**. PFA-fixed, coronal section stained with DAPI showing a recovered GRIN lens track in the dPAG (top, scale bar = 1 mm) and a zoom-in on the imaged region of interest with GCaMP6s-expressing VGAT^+^ neurons (bottom, scale bar = 100 µm, dashed line = approximate imaging plane). **B**. Placements of recovered GRIN lenses along the rostrocaudal axis of the PAG (N = 7 out of 8 experiments). The images in A are from animal BF118.2.

**Figure S5.**
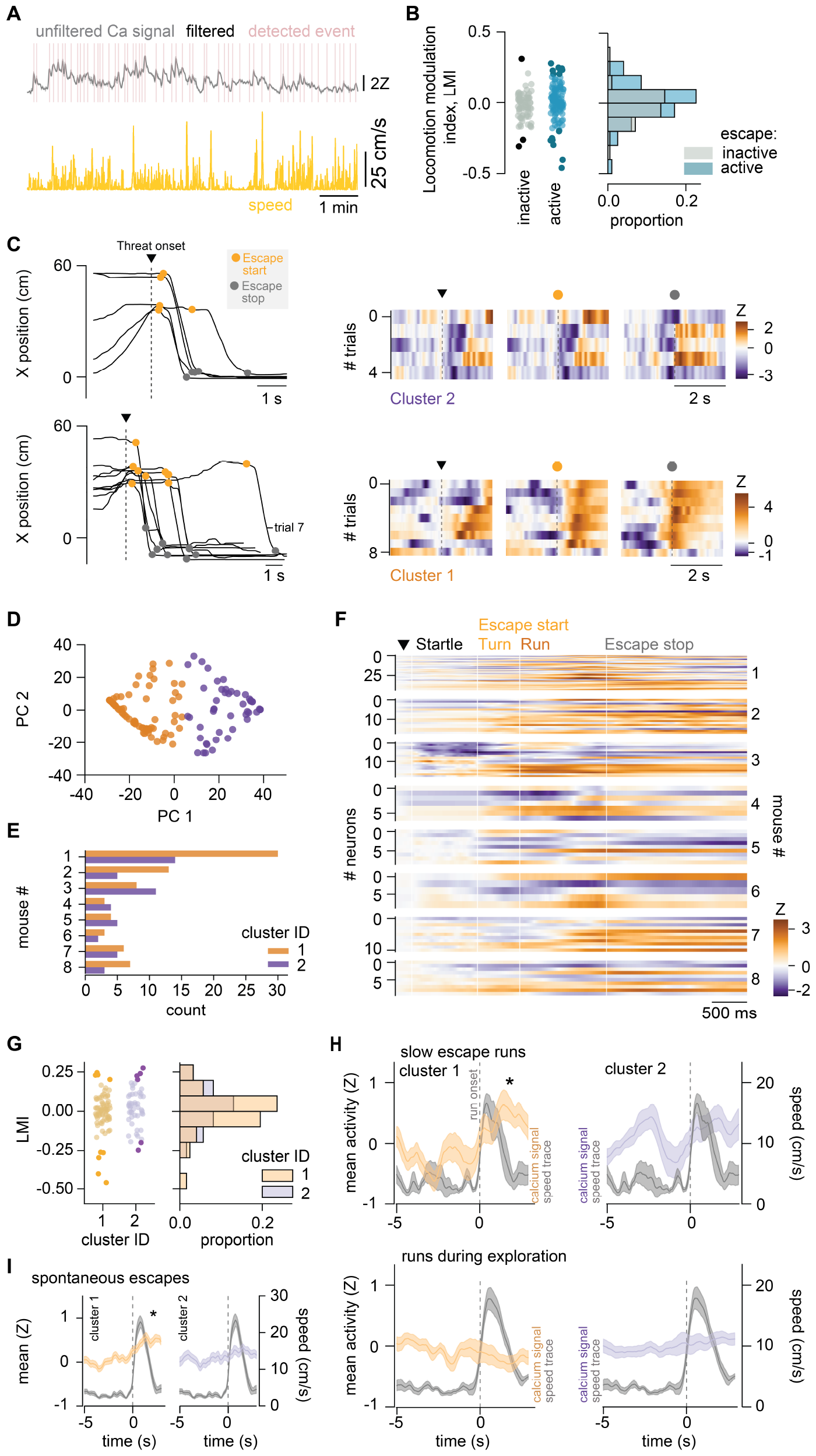
Analysis of calcium activity during exploration and spontaneous escapes and further examples of escape-active neurons. **A**. Single-neuron calcium signal (top) and animal’s speed (bottom) over time, showing calcium events (pink lines) detected using the filtered signal (black line). **B**. Distribution of cells’ locomotion modulation index of activity during exploration for populations that are inactive and active at escape. Dark coloured dots denote significantly modulated cells, determined by bootstrapping with resampling (95 % confidence interval significantly different from zero). Escape-active population contains a larger fraction of significantly modulated cells (8/123 and 7/123 cells, positive and negatively-modulated respectively) than the inactive population (1/76 and 2/76). **C**. Each row shows (left panel) all escape trials of one session of one animal, showing animal’s position along the long-axis of the rectangular arena (shelter entrance at X = 0 cm) over time, aligned to stimulus onset. Right: Heatmap of the calcium signal of one example neuron during the escape trials shown on the left, with the peak calcium signal aligned to threat-onset, escape onset (turn) and escape stop. **D**. PC scores for mean time-warped signal during escape for each neuron, coloured by K-means cluster ID (n = 123 neurons, N = 8 animals). **E**. Histogram of neurons assigned to clusters in G for each mouse. **F**. Mean time-warped signal for each neuron (same data as in Fig. 2G) for each mouse. **G**. Distribution of cells’ locomotion modulation index of activity for the two escape-active clusters. Dark coloured dots denote significantly modulated cells, determined by bootstrapping with resampling (95 % confidence interval significantly different from zero). Both clusters contain similar fractions of positively and negatively modulated cells (cluster 1, 4/74 and 5/74 cells; cluster 2, (4/49 and 2/49), positive and negative modulation index respectively), and the means of the cluster distributions are not significantly different (P = 0.08, Mann-Whitney test). **H**. Mean cluster population activity and animal speed trace for escape trials with slow peak run speed (top; mean peak speed 20.4 ± 1.9 cm/s) and fast running events (i.e., darting) during exploration by the same mice (bottom; mean peak speed, 20.2 ± 1.3 cm/s), aligned to the onset of running. Cluster 1 activity rises significantly during escape running but not exploratory running, while cluster 2 activity changes are not detected in either behaviour (*cluster 1, slow escapes*, mean signal -0.18 ± 0.20 Z and 0.39 ± 0.17 Z in the 2 s before and after run onset respectively, two-tailed paired t-test, P = 0.039, n = 18 neurons, 15 trials, N = 4 mice; *cluster 2, slow escapes*, mean signal -0.14 ± 0.21 Z before and 0.3 ± 0.16 Z after, two-tailed paired t-test, P = 0.11, n = 19 neurons, 15 trials, N = 4 mice; *cluster 1, runs during exploration*, mean signal -0.12 ± 0.12 Z before and -0.20 ± 0.13 Z after, two-tailed paired t-test, P = 0.66; n = 21 neurons, 8 trials, N = 4 mice; *cluster 2, runs during exploration*, mean signal -0.02 ± 0.11 Z before and 0.06 ± 0.11 Z after, two-tailed paired t-test, P = 0.72; n = 23 neurons, 8 trials, N = 4 mice). **I**. Mean cluster population activity and animal speed trace aligned to the onset of spontaneous escapes to the shelter in the absence of sensory stimulation (cluster 1, mean signal 0.01 ± 0.07 Z and 0.42 ± 0.07 Z in the 2 s before and after escape onset respectively, two-tailed t-test p < 0.0025, n = 61 neurons, 14 trials, N = 7 mice; cluster 2, mean signal 0.10 ± 0.09 Z before and 0.24 ± 0.08 Z after, two-tailed t-test p = 0.26, n = 44 neurons, 13 trials, N = 7 mice).

**Figure S6.**
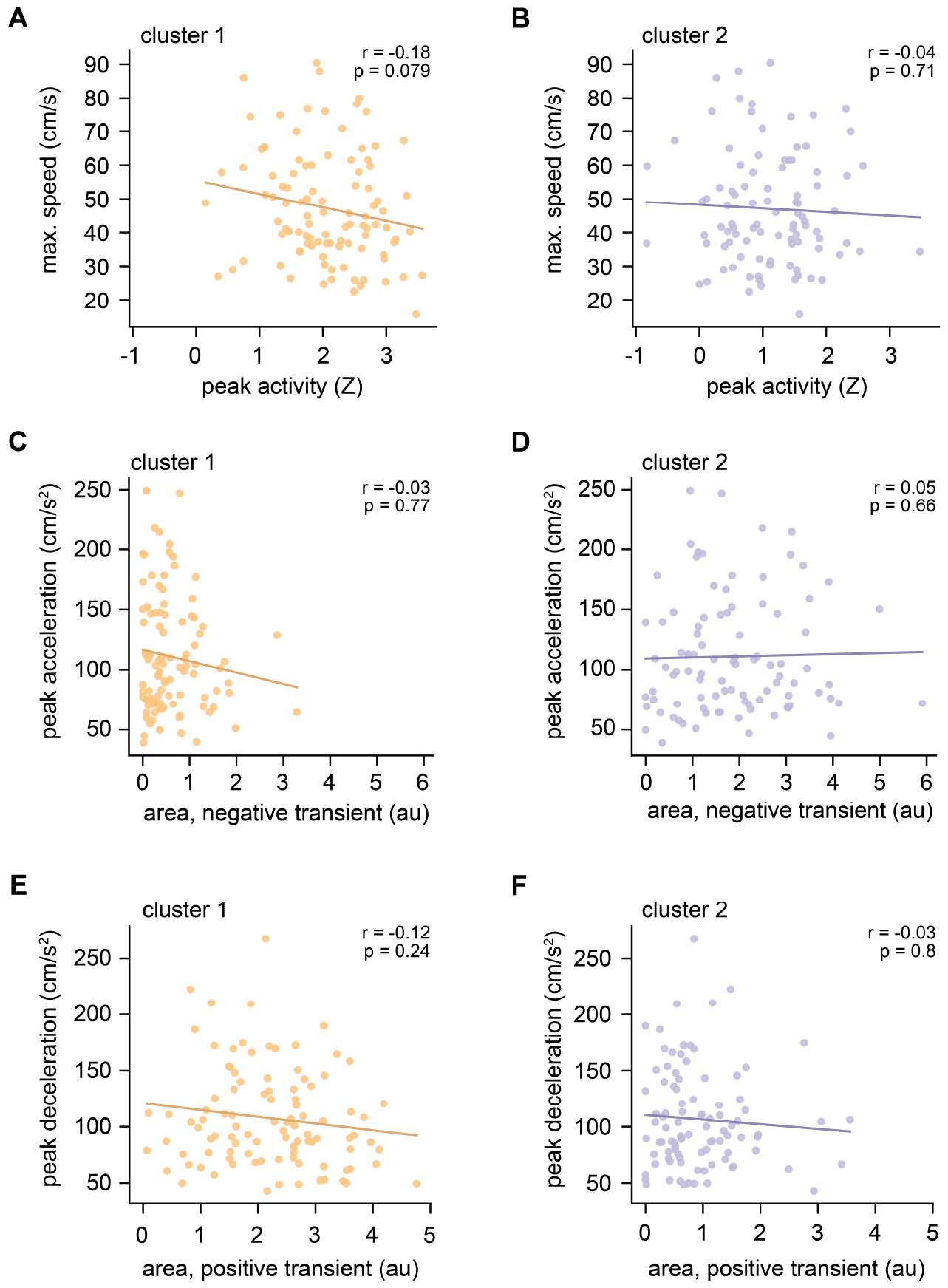
Population activity during escape is not significantly correlated with metrics of escape vigour. **A**. and **B**. Single trial mean peak Z-scored calcium activity during escape for cluster 1 neurons (**A**.) and 2 (**B**.) against maximum speed of the escape trial. **C**. and **D**. Area of absolute negative Z-scored calcium activity during escape for cluster 1 (**C**.) and 2 (**D**.), against peak escape acceleration from escape onset to time of maximum speed during escape. **E**. and **F**. Area of positive Z-scored calcium activity for cluster 1 (**E**.) and 2 (**F**.) against peak deceleration from time of maximum speed to escape termination. N = 100 trials, 8 animals; Spearman’s r and p-values are indicated for each measure in the respective panel.

**Figure S7.**
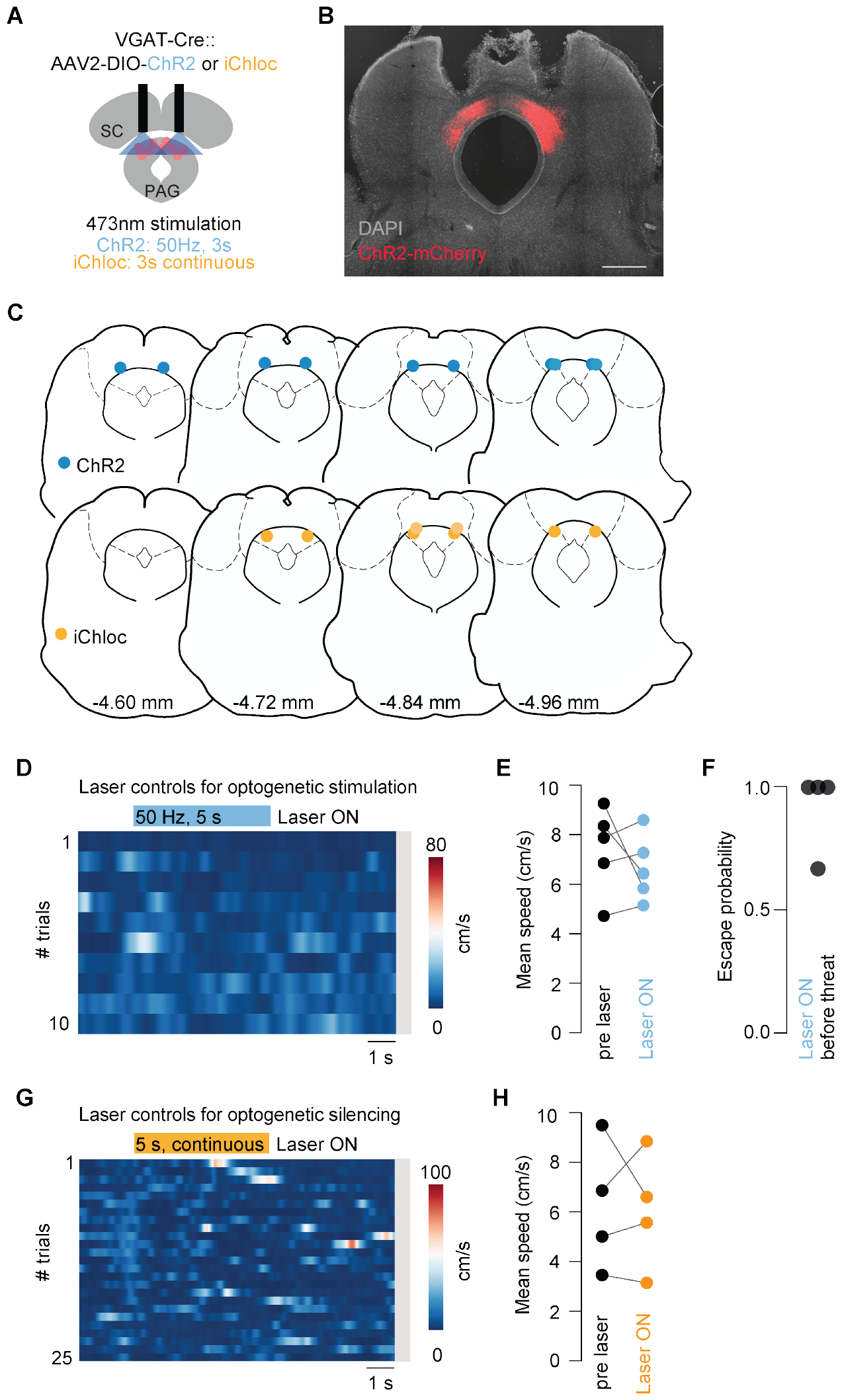
Optogenetic manipulation of VGAT^+^ dPAG neurons does not alter speed of movements during exploratory behaviour. **A**. Schematic of the experimental design and optogenetic stimulation protocols (ChR2: 50 Hz, 3 s; iChloC: continuous, 3 s). **B**. Coronal slice stained with DAPI (blue) with bilateral expression of AAV2/2-DIO-ChR2-mCherry (red) localised to the dPAG and optic fibre tracks. Scale bar = 1 mm. **C**. Bilateral optic fibre placements along the rostrocaudal axis of the dPAG are shown for ChR2 (blue, top; N = 5 mice) and iChloC experiments (orange, bottom; N = 4 mice). Rostrocaudal position from Bregma, based on Paxinos and Franklin. **D**. Example speed raster plot during optogenetic stimulation of VGAT^+^ dPAG neurons during exploratory movement. Grey bar on the right side of the histogram indicates no escapes. Laser duration is shown in blue. **E**. Summary graph of mean speed before and during laser stimulation (ChR2; N = 5 mice, n = 92 trials). **F**. Animals escape to threat stimuli when optogenetic stimulation of VGAT^+^ dPAG neurons precedes and ends immediately before threat stimulation (N = 4 mice, P = 0.44, Mann-Whitney test between probability of escape when laser precedes vs no laser condition). **G**. The same as D, for optogenetic inhibition. Laser duration shown in orange. **H**. Mean speed before and during laser stimulation for optogenetic inhibition (iChloC; N = 4 mice, n = 108 trials).

**Figure S8.**
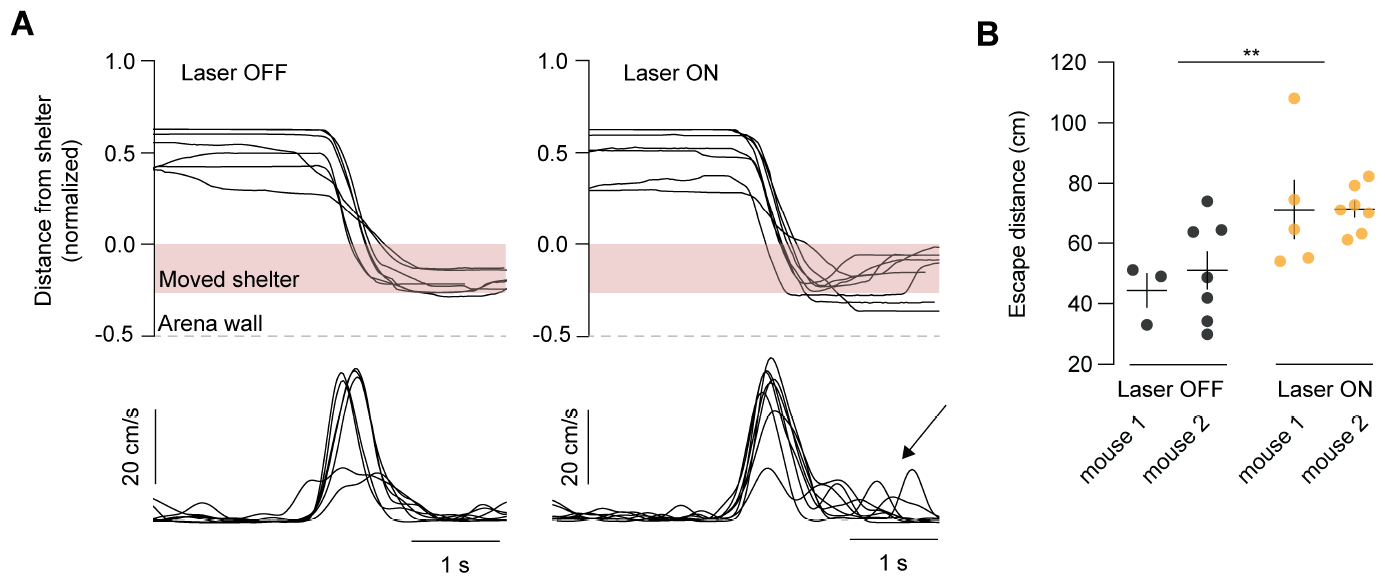
Optogenetic inhibition of VGAT^+^ neurons during escape increases total escape distance. **A**. Left: Moving the shelter location in the arena away from the arena wall leads to sensory-evoked escapes to the new shelter location (indicated in light red). Control trials (Laser OFF) of one mouse during escape to shelter with normalised tracking coordinates over time (top panel) and the corresponding speed traces (bottom panel). Right: Same as A, for Laser ON trials with light-activation of iChloC after escape onset. The same animal displays continued movement after reaching the shelter (black arrow). **B**. Summary plot of the total escape distance for Laser OFF trials (10 trials) and Laser ON trials (13 trials) showing a significant increase during light-activation of iChloC (N = 2 mice). Graph is showing the mean ± s.e.m and individual trials, separated by mouse.

## Notes

### Competing Interest Statement

The authors have declared no competing interest.

### Summary of Updates

Updated analysis and supplemental videos for optogenetic experiments added.

